# Crosstalk between plasma membrane and *Staphylococcus* α-hemolysin during oligomerization

**DOI:** 10.1101/2024.05.11.593496

**Authors:** Arnab Chatterjee, Anupam Roy, Thejas Sathees, Debajyoti Chakraborty, Partho Pratim Das, Bapan Mondal, Prithiv Kishore, Bartika Ghoshal, Siddharth Jhunjhunwala, Mahipal Ganji, Somnath Dutta

**Author notes:** Equally contributed (Arnab Chatterjee, Anupam Roy). Corresponding Author: Somnath Dutta.

## Abstract

The infectious microbe *Staphylococcus aureus* releases an array of cytotoxic pore-forming toxins (PFTs) that severely damage the cell membrane during bacterial infection. However, the interaction interfaces between the host cell and toxin were merely explored. Herein, we monitored the active oligomeric states facilitated membrane disruption processes such as lysis, and protrusion in the plasma membrane and lipid membrane. Furthermore, necrosis was triggered in the neutrophil-like cells upon synergistic binding and oligomerization of the monomeric α-HL. Additionally, we solved RBC membrane stabilized structure of different conformational states of this β-PFT using a single-particle cryo-EM. We further confirmed that internal membrane fluidity was the deterministic factor associated with the formation of intermediate pre-pores, heptameric pore-like, and complete pore species. Together, this is the first study to unveil the structure-function analysis of pre-pore to pore transition of any small β-PFT during its crosstalk with the cell.

**Highlights:**

1. α-HL promotes necrosis in HL60 cells and lysis of shorter lipid bilayer region.
2. Cryo-EM of small PFT in the cellular environment.
3. Structural characterization of heptameric pore, pore-like, and pre-pore complex in the presence of RBCs.
4. Bilayer phase behavior (Ld/Lo) governs different conformational and geometrical variants of α-HL.

## Introduction

The host cell plasma membrane, a complex heterogeneous mixture of protein and lipid, provides cellular integrity and indirectly governs several essential intracellular functions. This plasma membrane facilitates several fundamental cellular processes, like solute and metabolite transport, ionic balancing, intracellular signaling, and cell-cell signal transduction, by the membrane proteins adhered to plasma membranes ^1–3^. Thus, to hijack the host cell, the plasma membrane becomes an attractive target for several invading pathogens and their virulence factors^4–6^. Indeed, several deadly pathogens primarily arrest the host immune system and thereby escape the intrinsic cellular defense mechanisms^4,5,7^. Pore-forming proteins/toxins (PFPs/PFTs) are one of the significant classes of virulence factors, called exotoxins, secreted by a wide range of pathogens, which essentially target the host cell plasma membrane and tether the membrane-adhered receptors to promote inflammatory responses^8,9^. Alpha-hemolysin (α-HL/Hla) is one of the most abundant PFTs released by several opportunistic pathogenic strains of *S. aureus* that targeted the host immune cells^10–13^. The water-soluble, monomeric PFTs undergo several conformational changes upon binding to the host cell membrane and form an oligomeric complex with transmembrane pores^4–6^. Although how do these several oligomeric states of PFTs precisely relate to their cytotoxic nature remains unanswered. However, upon encountering such invader PFTs, host cell undergoes the membrane-mediated evagination and invagination as a part of cellular defense mechanisms^14,15^. These processes are involved in pore-induced increase of intracellular Ca^2+^ influx and followed-up processes, including the endocytosis of the pore-invaded regions and fission into several PFT-enriched micro-vesicles^15–17^. This Ca^2+^ influx was significantly high during the interactions between host membrane and large pore forming toxins, which accelerated the plasma membrane alteration^14,16^. Furthermore, sphingomyelinase mediated lysis of sphingomyelin (SM) into phosphocholine and ceramide facilitated such endocytosis process^3,14,15,18^. Nevertheless, the pore-forming toxin can promote bilayer re-organization, which further facilitates such defensive mechanisms, is still unanswered. Though the Cholesterol Dependent Cytolysin (CDC) and listeriolysin O (LLO) family toxins formed large membrane-embedded holes, and as it shared an extensive protein-lipid surface, the bilayer re-rearrangement was more feasible compared to small oligomeric PFTs like α-HL^4,19^. Toxin induced membrane blebbing and cellular protrusions were common phenomenon for CDC family toxins, like LLO, PFO, SLO, ALO, SLY, and LLY, which formed large ring-like complexes having pore diameters ranging from 25-50 nm^14,20^. However, these membrane-assisted blebbing and cellular protrusions were not reported for the small α-HL pore complex. Experimental evidence to find small PFTs-induced bilayer re-modulation is quite challenging. Moreover, a more difficult target is a structural understanding of transient intermediate states and their remodeled lipid-protein interfaces at atomic/near-atomic resolution.

This current study proceeded with the investigation of the impacts of one of the small PFTs: α-HL (a β-PFTs from *S. aureus*) on different cellular environments, like RBC, innate immune cells, to identify the membrane deformation including vesicle blebs, fusion, lysis, at a nanomolar scale. Additionally, interactions of PFTs with several artificial lipid compositions, like phosphatidylcholine (PC), PC-lipids with cholesterol, and sphingomyelin were examined to understand lipid-PFT interactions. Additionally, we also provided a detailed cellular response, especially toxin-induced morphological change in different host cell plasma membranes and lipid vesicles. We resolved several atomic resolution structures of complete pores, pore-like, and membrane anchored pre-pore conformations of α-HL using cryo-electron microscopy (cryo-EM). Furthermore, our study is the first experimental evidence where we employ fluorescence-based imagining, biophysical, biochemical approaches, cellular-level approaches, and cryo-EM-based structural studies to address the pore-forming mechanism and associated membrane re-modulation.

## Results

The plasma membrane is the protective shield of the cell and other integral compartments and organelles within cells, composed of the complex architecture of a lipoprotein-lipid membrane bilayer with an overall width of 30-40 Å^1–3^. The local protein-lipid interactions were generally governed by the cooperation of membrane-inserted protein folding and the lipid bi-layer modulation^21,22^. However, various bacterial virulence factors, especially the pore-forming class of proteins, can destabilize the lipid bilayer integrity. Herein, to unfold α-HL and plasma membrane crosstalk, we depicted the cellular responses of innate immune cells and RBCs against sub-nanomolar and micromolar concentrations of α-HL.

### Characterization of α-HL-mediated effect on the host cell membrane

To unveil the α-HL-plasma membrane crosstalk, we purified the water-soluble recombinant toxin α-HL using Ni-NTA-based immobilized metal affinity chromatography (IMAC) followed by size exclusion chromatography (SEC). The purity of the isolated recombinant protein was analyzed by SDS-PAGE and SEC profile ***(Figure 1A, S1A, B)***. MALDI data suggested the purified monomeric toxin having a molecular weight of ~35kDa ***(Figure S1D)*** and isolated monomeric α-HL showed a thermal stability (T_m_) upto 50°C ***(Figure S1E)***. The functional activity of the purified toxin was confirmed using the haemolytic assay, which showed concentration-dependent (0.01-0.1 μM) damage of the rabbit erythrocyte plasma membrane ***(Figure 1B)***. Further, we wanted to explore the toxin-mediated effect on innate immune cells as well as on RBCs. The SDS-PAGE analysis depicted a higher-order oligomeric state of α-HL upon incubation with HL-60 and washed erythrocyte cells ***(Figure S1C)***. To understand oligomer-induced membrane damage, a DNA-binding dye, 4’,6-diamidino-2-phenylindole (DAPI) was introduced along with the toxin (50 nM) in the growth medium of RAW264.7 and HL-60, which resulted in labelling of the nucleus of macrophage cells after α-HL treatment ***(Figure 1C, S2A)***. HL-60 cells initially stained three lobes of the nucleus because of DAPI binding to nucleus; however, after 5 minutes, intensity of DAPI is significantly reduced from HL-60 ***(Figure 1C)***. This might be possible that the nucleus of the HL-60 was dispersed from the HL-60 cells over the time due to membrane rupture. However, DNA intensity was intact inside macrophage cells over the same period of time after toxin attack (***Figure 1C, upper right-hand panel, Figure S2A***). We observed a slight decrease in Nile red intensity from the surface ***(Figure 1D)*** of HL-60, which could be due to untethering of the lipid molecules after pore formation. Additionally, our study showed shrinkage of HL-60 cells and protrusion of small vesicles from the plasma membrane ***(Figure S2B, C)***. Additionally, we noticed the clumping of HL-60 cells and a decrease in average cell size by 2µm after 10 minutes of α-HL treatment ***(Figure S2C, D)***. To determine the cell-death pathway induced by the purified toxin in a quantitative manner, flow cytometry of toxin-treated cells was performed using Annexin V and Ethidium homodimer III (EthD-III) fluorophores. At 10 nM toxin concentrations, HL-60 cells survived, whereas, after incubating with higher concentrations (100nM, and 1µM) cell populations shifted toward the necrosis-mediated cell-death pathway ***(Figure 1E, S2E)***. HL-60 cells also showed membrane deformation under the transmission electron microscope (TEM) and scanning electron microscope (SEM) ***(Figure S3A-C)***.

**Figure 1:**
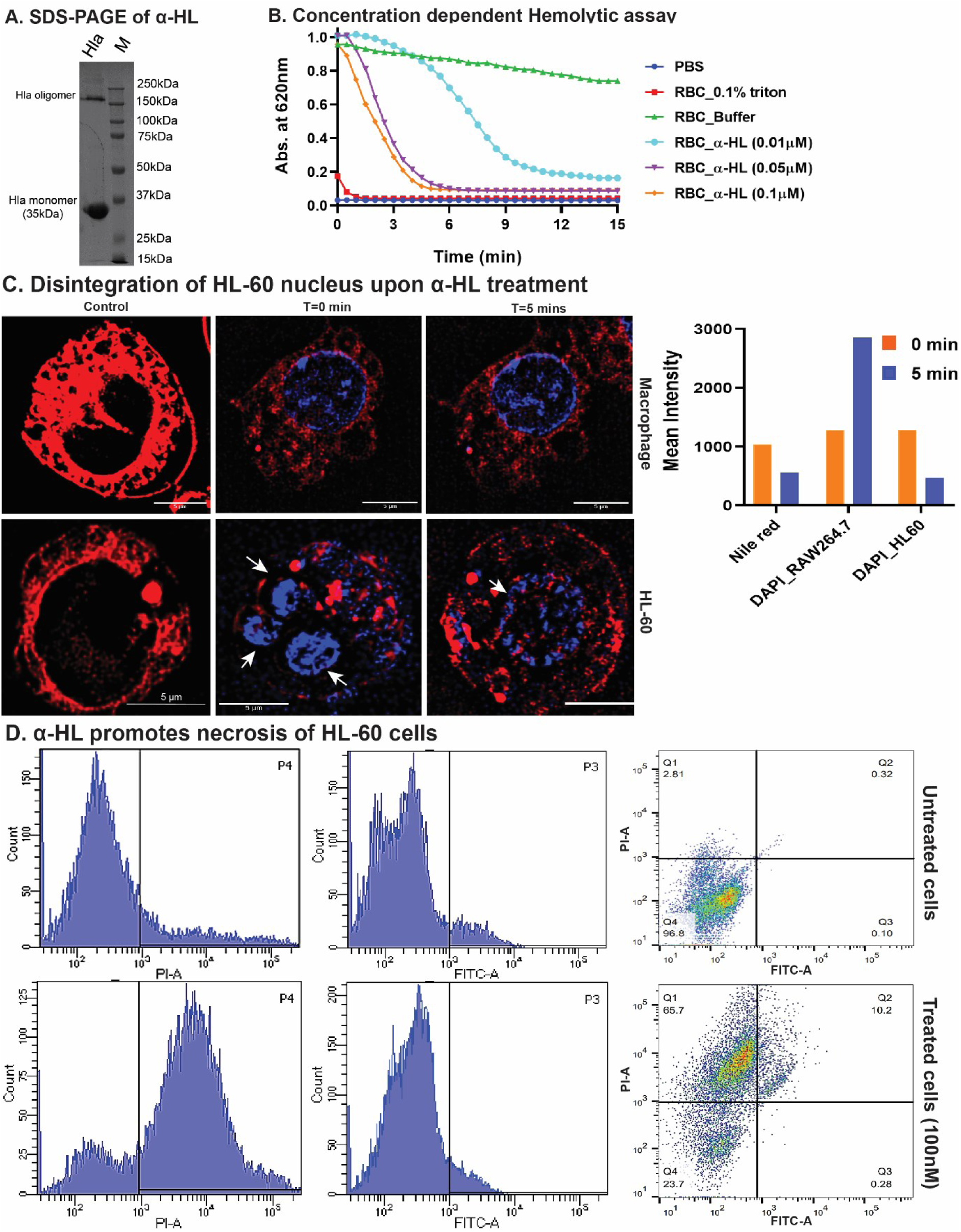
Small PFT (α-HL)-mediated effects on the host cell. **A.** SDS-PAGE of SEC purified α-HL monomer. **B.** Hemolytic assay of rabbit erythrocytes (10%) was performed with different concentrations of α-HL toxin. **C.** Macrophage & HL-60 cells were incubated with α-HL (50nM) respectively in the presence of nucleus staining dye, 4′,6-diamidino-2-phenylindole (DAPI, λ_em_= 460 nm) & membrane staining dye Nile red, (λ_em_= 570 nm). **D.** The relative change in fluorescent intensity in the Nile red intensity in RAW264.7 cells and the change in DAPI intensity for both the cells over time were plotted (left to right). Orange-red and blue-colored bar graphs represented before and after toxin incubation respectively. **E**. Flow cytometry analysis of untreated and 100nM toxin-treated HL-60 cells stained with annexin V-FITC and EthD-III. Compared to the untreated cells, the toxin-treated cells started becoming necrotic as evident by the shift towards EthD-III gate.

Additionally, we observed normal discocyte-shaped RBC, schistocytes (fragmented) ***(Movie 4, 5)***, different stages of sphero-echinocyte (tear-shaped protrusion) ***(Movie 6, 7)***, and upon α-HL treatment on freshly washed RBCs ***(Figure 2A***). Negatively stained TEM images showed that the toxin-treated RBC membrane was severely distorted and the leakage of the cytosolic material through the ruptured RBC membrane, which was completely absent in control RBC ***(Figure 2B)***. Thus, these TEM data suggested that toxin induced significant damage of RBC plasma membrane.

**Figure 2:**
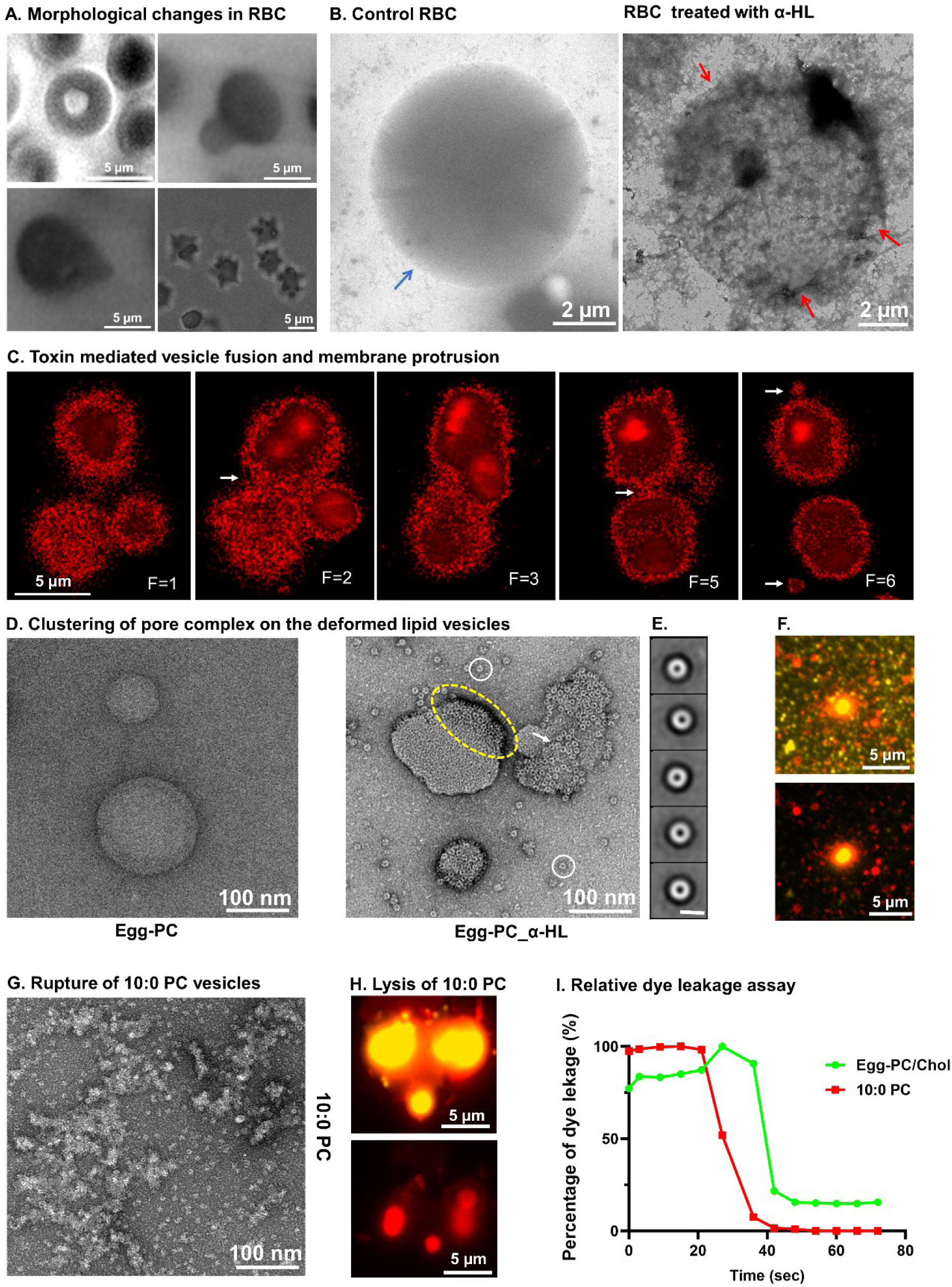
Impacts of toxin on RBC plasma membrane and other composites lipid membranes. **A.** Change of RBC shape was captured using TIRF, and confocal microscopy. **B.** A partial lysed rabbit erythrocyte plasma membrane and leakages of intracellular material (red arrowhead) after 5 minutes α-HL (100nM) toxin treatment. **C.** Snapshots of different fates of liposomes after toxin treatment including vesicle-vesicle fusions and protrusion of lipid bodies (white arrows). **D.** The NS-TEM data of α-HL added to egg-PC/cholesterol liposomes showed the formation of distinct pore complexes on the lipid membrane (white arrowhead, yellow encircled). **E.** NS-TEM 2D class averages of oligomers (scale bar, 10nm). **F.** Single-molecule imaging shows leakage of intra-vesicular dye (rhodamine dye, λ_em_= 570 nm) after α-HL (10nM) treatment on NBD tagged (λ_em_= 540 nm) eggPC/Chol LUVs. **G.** NS-TEM analysis showed the complete rupture of 10:0 PC LUVs after 30 minutes of toxin (50nM) treatment. **H.** TIRFM imaging confirmed the lysis of fluorescent NBD-labelled liposome and the subsequent leakage of encapsulated rhodamine dye while passing of α-HL in the microfluidic channel. **I.** Rhodamine B (λ_ex_= 540 nm, λ_em_= 570 nm) leakage kinetics upon toxin treatment on 10:0 PC LUVs and egg-PC LUVs.

### α-HL mediated alterations of lipid membranes

As the lipid binding motifs of small PFTs primarily interact with membranes to form functionally active oligomers^23^, we were curious to understand the mechanism of toxin-induced alterations of the artificial lipid membrane. Previous studies reported that phosphatidylcholine (PC) favored the oligomerization of toxins^23^; thus, the egg-PC-containing liposomes were used to check the toxin-mediated oligomerization effect of α-HL on the lipid membrane. Additionally, to monitor the impact of α-HL on liposomes, fluorescently labeled phosphatidyl ethanolamine (Rhod-PE) was incorporated with egg-PC and cholesterol-mixed liposomes. Upon α-HL treatment, observe a few vesicles fusions process and small micro-vesicles thrown out as protruding bodies from egg PC/Chol vesicles, which could be as an outcome of membrane stabilization ***(Figure 2C; S4A; S5A; Movie 9-11)***. The fluorescently labeled protein co-localized with Rhodamine-tagged liposomes confirmed the presence of the toxins on the egg PC/Chol liposomes (***Figure S4C***).

The outer leaflet of cell membrane composites of different phosphatidylcholine lipid chain lengths^24^. Thus, we prepared liposomes of various phosphatidylcholine lipid chain lengths (10-Carbon PC to 18-Carbon PC) to monitor the role of lipid chain length in oligomerization of toxin and membrane damage. The overall shape of the membrane was intact in the case of egg-PC after toxin treatment ***(Figure 2D, F; S5A)***, though α-HL rendered 10:0 PC liposomes into complete ruptured conditions (***Figure 2G, H; S5A)***. Similar oligomeric pore-like species were also observed on the disrupted lipid membrane, along with some higher-order assemblies of heptameric geometry ***(Figure S5C)***. However, liposome rupture was further reduced for longer chain lengths of liposome (14:0 PC and 18:1 PC), which was completely abolished in presence of 16:0-18:1 PC (***Figure 2D; S5B)***. This provided a preliminary idea that small pore-forming toxins can significantly impact the cell membrane, where hydrophobic chain length of lipid was comparatively shorter. We employed single-molecule imaging approach to elucidate the real-time changes in the lipid vesicles fluorescence intensity using total internal reflection fluorescence (TIRF) microscopy. Large Unilamellar Vesicles (LUVs) composed of 10:0 PC with a fraction of biotinylated and fluorescently tagged PE were loaded with a fluorescent dye Rhodamine B. These biotinylated liposomes were then immobilized on streptavidin coated PEG-passivated coverslips. α-HL was flowed in at a fixed flow rate onto the immobilized egg-PC and 10:0 PC liposomes separately at 10 nM concentration. The liposome surface was visualized by excitation using a 488 nm laser, and the internal rhodamine dye using a 561 nm laser. The changes happening on both channels were recorded simultaneously. Buffer wash did not release the liposome encapsulating dye ***(Figure S6D-G; Movie 12, 13)***. We observed that the immobilized liposomes released the internal dye content at around 40 seconds from both types of lipid vesicles ***(Figure 2F, H, I; Movie 14, 15)***. These results suggested that the impact of PFTs on the lipid bilayer depended on the properties of the membrane. However, the mechanism of vesicle disruption by PFT remained elusive. Thus, we opted for cryo-EM-based structural approach to understand lipid binding regions and associated conformational states of α-HL that occurred during membrane perforation process.

### The structural and molecular form of α-HL pore complex in RBC plasma membrane

To understand the molecular insights of the cytolytic form of α-HL with the plasma membrane at the near-atomic resolution, we employed the single-particle cryogenic electron microscopy (cryo-EM) approach to characterize 3D structure of α-HL in the presence of real cellular environment. The cryo-EM micrographs of toxin-treated lysed innate immune cells and RBCs both showed the formation of α-HL oligomers ***(Figure S7A-C)***. However, due to leakage of large intracellular components from innate immune cells, the RBC plasma membrane has been considered as an ideal model cellular environment for further cryo-EM analysis (***Figure S8***). From our cryo-EM study, we observed ring-shaped proteins assembled on the surface of the RBC plasma membrane and well-dispersed oligomeric species detached from the RBC membrane ***(Figure 3A)***, where high-resolution structural features were clearly visible in reference-free 2D class averages (side and top view). A followed-up cryo-EM reconstruction of this cytolytic protein complex provided the first solution states cryo-EM reconstruction in the presence of RBC plasma membrane-embedded α-HL pore complex at a 3.5Å global resolution ***(Figure 3B)***. The cytolytic β-barrel domain was composed of fourteen identical transmembranes (TMs) segments (an antiparallel β-strand forming segment Y112-T125 and G135-T145) as depicted in cryo-EM surface representation (***Figure 3B, C***). The transmembrane (TMs) heptameric α-HL complex formed a 35.5 Å (from O_η_ atom of C_α_ Y112 to C_α_-I132) long symmetrical channel with an average pore diameter of 2.57 nm (C_α_ N121 to C_α_ A138) (***Figure 3C***). The large extracellular domain of this pore complex consisted of a β-sandwich cap and a partly unfolded rim domain (***Figure 3B, C; S9B*)**. PC head group was fitted in the pore lined lipid densities present near the upper β-barrel region (***Figure 3D; S9C***), which was the first evidence of interaction of lipid head groups with the β-barrel PFTs (***Figure S9D***). Such amphipathic-aromatic lipid head group anchoring residues were also shown to stabilize the polar interfaces of lipid bilayer for other β-PFTs, including anthrax toxin, and lysenin^25,26^. However, in the interior of the lipid bilayer shell, the TMs hydrophobic residues (L116, Y118, F120, V140 & I142) were extensively interacting with the hydrophobic lipid wall (***Figure S9E***). On the other hand. the inner architecture (cross-section) of the TMs channel and extracellular pore-vestibule were predominantly polar **(*Figure S9F, G*)**. The variable amphipathic features of the TMs segment α-HL pore suggested having more hydrophobic and aromatic residues in the top part of the lytic domain. Whereas the lower half of the lytic domain contains less polar and flexible amino acids (G122, V124, G126, G134, I136, A138) ***(Figure S9E)***. The relative thermal fluctuations were quite large in the TMs region compared to other segments of the pore complex, especially in the bottom part of the lytic domain and cytosolic segment (***Figure S9H***). However, cryo-EM structural analysis of the oligomeric states showed an additional heptameric pre-pore complex where the density of TMs segment was entirely unresolved from our cryo-EM reconstituted map (***Figure S10A***).

**Figure 3:**
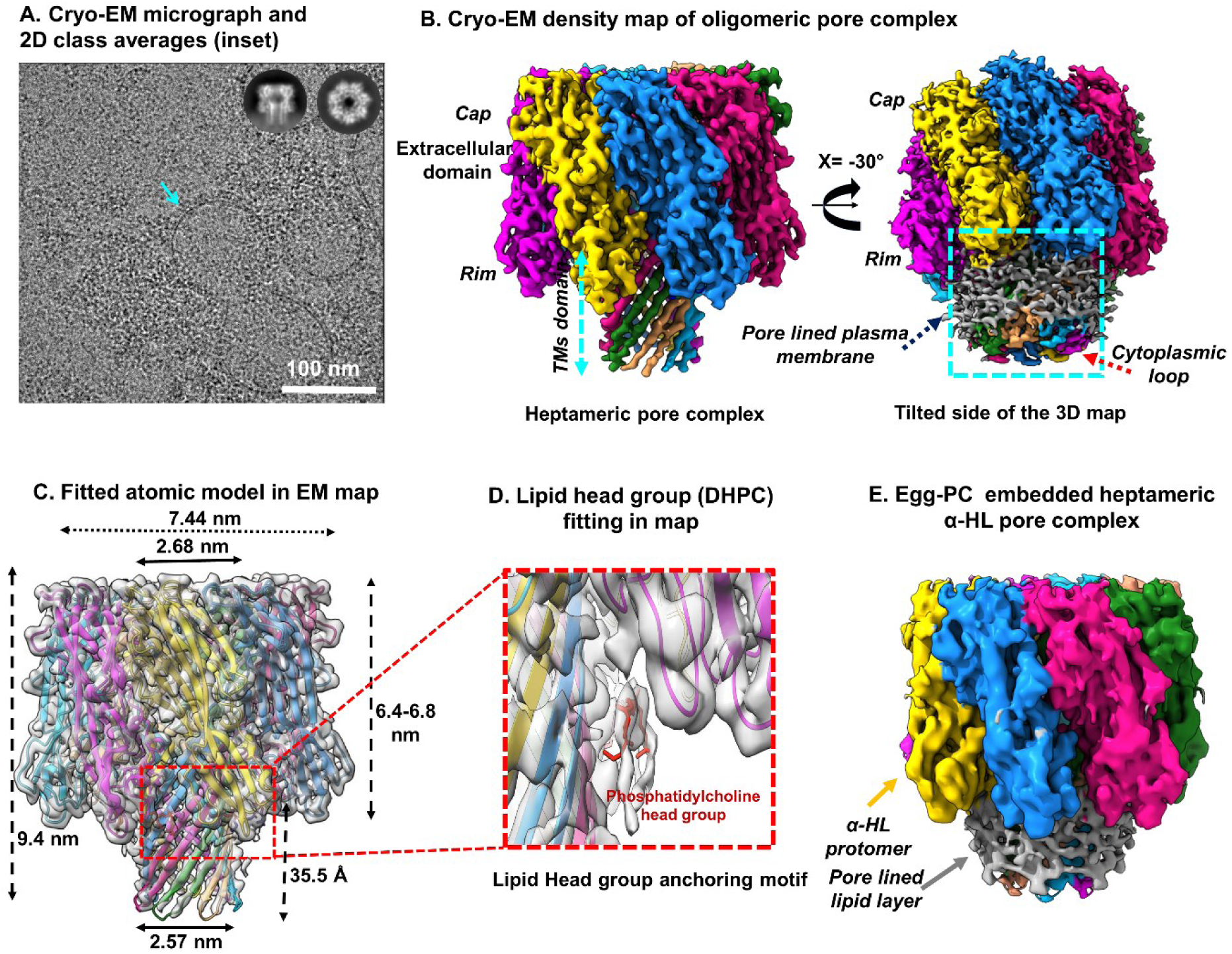
Cryo-EM analysis of pore formation of α-HL in the native cellular (RBC) environment. **A**. Representative cryo-EM micrograph showed the distribution of pore particles embedded (cyan arrow) on the RBC micro vesicles. 2D class averages (inset) showed the heptamer (top view) and mushroom-shaped complex (side view). **B**. 3D reconstruction of heptameric pore complex resolved at 3.5 Å global resolution in RBC membrane environment. The tilted lateral view (X= −30⁰) depicted the pore-lined lipid arrangement (cyan box) around the transmembrane (TM) channels (cyan arrow) (threshold:0.0116). **C.** The lateral view of a cartoon representation of atomic model fitted in cryo-EM density map of heptameric pore. **D.** Selected enlarged view showing the fitting of the atomic density head group of PC (ball & stick model; red). **E.** Cryo-EM 3D refined map of pore complex generated from egg-PC/Cholesterol model membrane.

A simultaneous co-existence of such pre-pore states (30%) with abundant pore complexes in the RBC plasma membrane could be an intermediate species that appeared as a dynamic process of pore formation (***Figure S10A***). Therefore, it might be possible that α-HL assembled as pre-pore and pore conformations in presence of RBC membrane, where ADAM10 receptor were present. However, our observations also suggested that α-HL had an oligomerization propensity towards the phosphocholine (PC) lipid bilayer ***(Figure S2D)***. This suggested that either receptor or physical/chemical property of lipid play a vital role in actual pre-pore and pore formation. To understand the role of the lipid in α-HL pore formation, α-HL was structurally characterized in the presence of liposomes, which constituted by single lipid component (egg-PC, 10-PC, 14-PC).

### The structural metamorphosis of α-HL pore complex in proteo-liposomal membranes and α-HL induced re-organization of the lipid bilayer

The NS-TEM and cryo-EM data suggested that α-HL was assembled as oligomeric complexes over LUVs surfaces (***Figure 2D, E; S11A***). Furthermore, we determined the cryo-EM structure of heptameric pore structure of α-HL in presence of 16:0-18:1 PC/Chol (***Figure 3E)***. Indeed, 38-40 Å lipid bilayer thickness of the egg-PC permitted the hydrophobic stabilization of the entire lytic β-barrel domain (36 Å) of the heptameric pore complex (***Figure S11A***).

Unlike egg-PC (16:0-18:1 PC), and 14:0 PC, a rapid lysis of 10:0 PC vesicles was observed during toxin-induced membrane perforated complex formation (***Figure S5A, B***). Thus, to understand lipid lysis vividly and the lipid-protein hydrophobic interactions, which might play a significant role in membrane damage, we further shifted our model membrane into the shortest acyl chain containing lipid vesicles composed of 10:0 PC (~ 26 Å thick)^27^. It would be interesting to use a model membrane composed of 10:0 PC, where the hydrophobic amino acids of β-barrel would have been exposed to the outer hydrophilic environment. However, our cryo-EM study and structural analysis of oligomeric species with 10:0 PC bilayer strongly suggested that 36 Å long hydrophobic β-barrel lytic domain of pore complex was significantly stabilized by 10:0 PC (***Figure 4A***). Surprisingly, the structural features of the pore complex identified from the plasma membrane and egg-PC/Chol composite model membranes resembled pore complexes isolated from lysed 10:0 PC vesicles (***Figure 3C; 4A***). Apart from pore line TMs stabilized lipid layer, we also identified an obscure fuzzy density located at the lipid recognition motif of the lower rim domain (***Figure 4A***). The abundance of amphipathic amino acids like W179, Y182, Y191, Q194, and R200 was close to such obscure lipid density at the lipid recognition domain (***Figure S12D***). However, W179 and R200 were initially identified as phosphocholine (PC) head group binding residues from the monomeric crystal structure of wild-type α-HL with phosphocholine^23^, which was correlated with our current study. From our investigation, we have predicted several amino acid residues like, aromatic residues W179, R200, and Y191 located at the lipid recognition motif and the lipid head group sensing amino acids Tyr148, Tyr112, and His144 detected at pre-stem lytic domain, which could be responsible for α-HL pore formation. However, due to the shorter acyl chain length, the bilayer length of 10:0 PC vesicles was shorter than egg-PC vesicles^27^. Cryo-EM analysis further calculated the intermediate oligomeric complex geometrically similar to the pre-pore states as observed in the plasma membrane of rabbit erythrocytes (***Figure 4B***). In comparison with the (50%) pore complex, the partly folded (45%) pre-pore complex contained a premature (9-12 Å) long non-TMs β-barrel located at the neck region of the stem domain. Interestingly, such (9-12) Å long twisted β-barrel stem segments (D108-Y112 and T145-P151) remained stabilized without lipid shell (***Figure 4B; S12B***). A non-lamellar (micelle) form of lipidemic arrangement had been identified in the pre-pore state that was located at the base of vestibular fenestration (***Figure 4B; S12E***). This blocked the vestibular channel and prevented the insertion of the lytic TMs pore. Our current results demonstrated that various lipid chain-lengths played an important role in pre-pore and pore formation. We were also curious to know how stability of lipid membrane affecting the pore formation. Therefore, our next strategy was to introduce the sphingomyelin in lipid membrane to visualize the membrane stability in the presence of α-HL.

**Figure 4:**
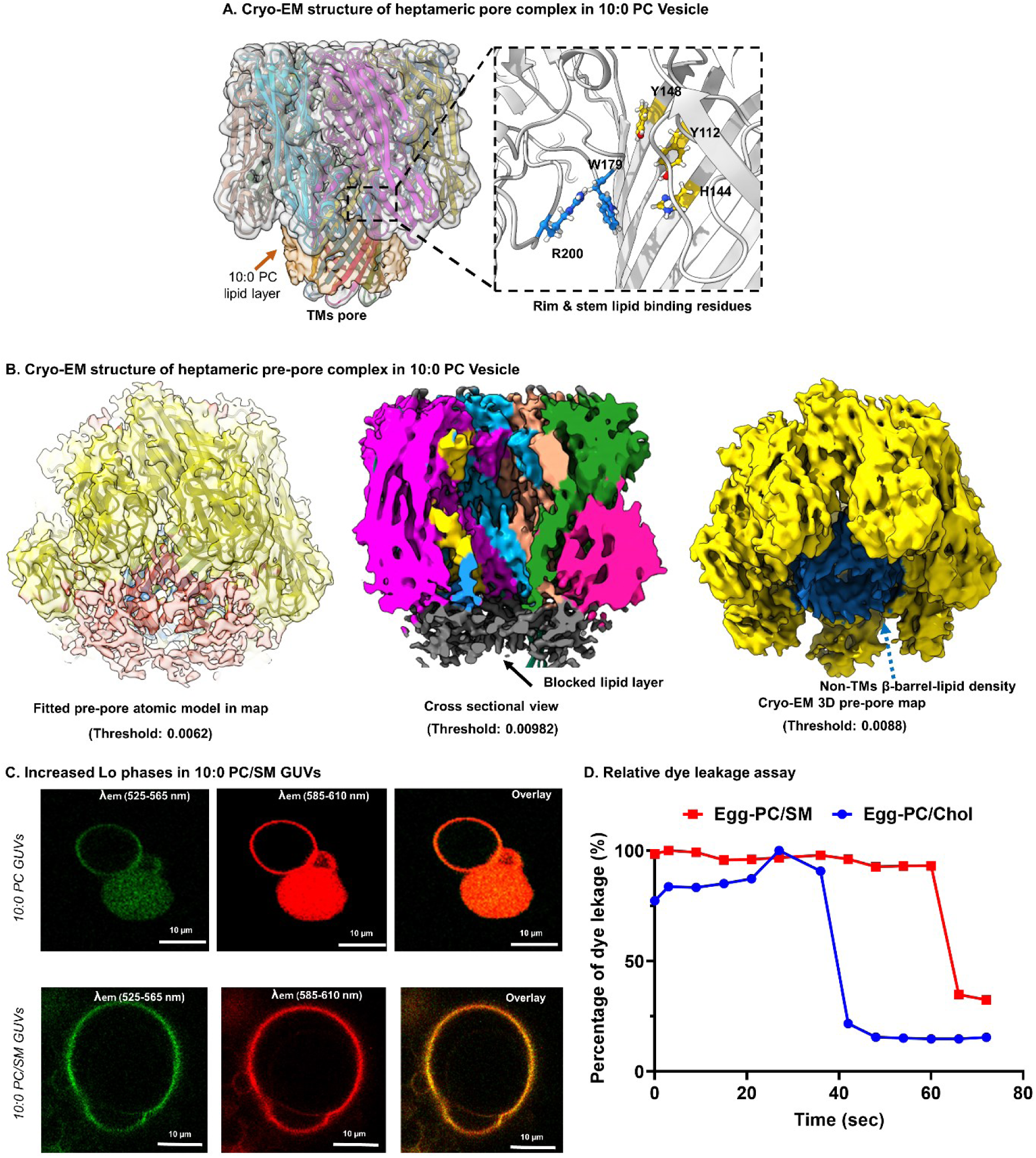
10:0 PC lipid layer upholds TMs stability despite of promoting heptameric pre-pore complex formation. **A.** The cryo-EM map of heptameric pore complex (gray) fitted with atomic model (cartoon). The 3.6 nm long lipid densities (light orange) surrounding the lytic TM channels. The enlarged view showed the residues (Y112, H144, Y148, W179, and R200) close to the lipid moieties located at the junction of the rim (blue) and the lytic stem domain (gold). **B.** Cryo-EM map (white) of heptameric pre-pore complex aligned with its atomic model (extracellular domain in yellow, non-TMs in blue) and extra lipid density in light orange. The cross-sectional view of the map (threshold 0.00982) having a thick lipid density (black) blocked the TMs pore vestibular channel. EM map (yellow; threshold 0.0088) sighting an extension of the non-TMs barrel (blue) located at fuzzy lipid density area in heptameric pre-pore state. **C.** Confocal images of 10:0 PC and 10:0 PC/SM in the presence of lipophilic membrane staining dye Nile red. SM incorporated 10:0 PC showed a relatively higher emission intensity of Nile red at 525-565 nm. **D.** Single molecule leakage assay of eggPC:SM vesicles with α-HL showing a slower lag phase kinetics compared to only eggPC:Chol vesicles.

Sphingomyelin, an essential lipid content located at the phosphocholine-enriched RBC plasma membrane, preserved the plasma membrane rigidity through firm adhesion with cytoskeletal proteins^28^. To unfold the role of sphingomyelin (SM) in the phosphocholine membrane bilayer, we treated monomeric α-HL toxin with 30% SM incorporated composite vesicles of phosphocholine (egg-PC and10:0 PC separately). We examined the liquid order phase of LUVs using a solvatochromic fluorescence dye (Nile Red) that showed high intense emission spectra at 585-610 nm in 10:0-PC/SM vesicles compared to egg-PC/Chol when excited at 488 nm (***Figure 4C***). In addition, sphingomyelin mixed lipid vesicles showed a delay in dye leakage as compared to the 10:0 PC and egg-PC lipid vesicles ***(Figure 4D; S6H)***. Further, cryo-EM analysis of α-HL toxin treatment on LUVs composed of 10:0 PC/SM resolved a distinct heptameric pre-pore and pore-like states (***Figure S12F, G; Figure S13***), which were structurally identical to previously predicted pre-pore and pore conformations. Previous reports and our observation suggested that the incorporation of SM into the PC bilayer increased the L_O_-rich lipid microdomain^3,14^, which lead to an incomplete pore formation. Therefore, we manipulated the phase behavior of the lipid bilayer (bilayer fluidity) physically without further incorporation of SM.

### The lipid phase behavior evolved as a prime factor controlling the α-HL pore maturation

We treated the pre-cooled egg-PC/Chol LUVs with monomeric α-HL to validate the hypothesis of pre-pore species formation with the increase in the ordered lipid. The pre-cooled egg-PC/Chol lipid vesicles were further reduced the dye leakage activity compared to the room temperature vesicles **(*Figure S6H; Movie 16*)**. Surprisingly, from the cryo-EM micrographs and 2D class averages of pre-cooled egg-PC/Chol incubated with a monomeric toxin, we observed the abundance of variable larger geometrical states (8-mer and 9-mer) along with heptameric particles ***(Figure S14A)***. However, we further proceed with pre-cooled rabbit RBC to investigate how the phase behavior of rabbit erythrocytes could modulate the haemolytic nature and formation of related cytotoxic forms of α-HL.

The haemolytic assay with pre-cooled rabbit erythrocytes at different toxin concentrations showed an increase in the lag phase of the haemolysis performed at reduced temperature ***(Figure 5A)***. Similar to egg-PC/SM composite vesicle, we observed an intense emission at 525-565nm range in the pre-cooled RBC membrane. This indicated the relatively compact packing of the RBC membrane component that led to forming a liquid-ordered phase of the plasma membrane ***(Figure 5B)***. Due to restricted membrane perforation, a pull of oligomeric states, including arc-shaped small oligomers and higher-ordered oligomers like distorted octamer, were detected, as shown in cryo-EM 2D class averages. However, unlike distorted octamer, we also observed 4-mer, 5-mer, and 6-mer formed arc-shaped oligomers that maintained a circumference identical to the heptameric states (***Figure 5D; Figure S14B***). In addition to a delayed and partly compromised haemolytic assay, the occurrences of mature heptameric pore species were not further identified from a detailed cryo-EM analysis ***(Figure S14B)***. The cryo-EM reconstruction was determined to have a 90% pre-pore and 10% pore-like heptameric complex apart from other intermediate oligomers ***(Figure S14B)***. However, the absence of the mature pore complex suggested that the pore-like heptameric states might be functionally active in pursuit (RBC) cell lysis. The pore-like states maintained similar backbone structural rearrangements with mature pore complex except for the unstructured bottom-half TMs β-barrel ***(Figure 5E, F)***. Additionally, the N-terminal stretch (A1-K8) interacted strongly with the inner part of the cap domain of two adjacent protomers ***(Figure 5G)***. The relative position of the N-terminal segment to its adjacent protomer in pore-like states suggested that the amino stretch (A1-K8) could further sterically trigger the pre-stem detachment from the inner surface of the cap domain during pore formation ***(Figure 5G).*** Furthermore, we identified the pre-pore state, devoid of TMs segments that could remain flexible (dynamic conformation) to displace the outer leaflet of the RBC plasma membrane. Thus, the lipid densities present at the bottom part of the rim domain consisted of both pre-stem, stem, and lipid moieties at the cytosolic vestibule region ***(Figure 6A)***. Cryo-EM results further suggested that the pre-pore complex was accompanied by the missing densities at β-turn segments of the extracellular upper cap domain, N-terminal loop (T9-V20). Those missing densities could be due to their high thermal backbone fluctuation. ***(Figure 6B, C)***. However, we could observe the density at the partially stabilized N-terminal segment (A1-K8) in the pre-pore states, which primarily controlled the pre-stem detachment process ***(Figure 6C)***. We performed a structural comparison with the protomers identified from pre-pore and mature pore complexes ***(Figure 6D-F)***. Detailed structural studies showed the inwards movement of β1 (K21-V26) and β2 (Y40-I44), located close to N-terminal of cap segments. Additionally, outer β-sandwich cap segments (Y282-I284) at β13 and (E289-T292) at β14 also followed an inwards movement ***(Figure 6D)***. Whereas the inner cap segments β5 (Q97-Y101), β10 (D227-M234) and the outer cap segments β9, (K164-F171), β4 (K85-L90) were also directly connected to the TMs stem domain followed an outwards left-handed movement. Whereas the β-turn segments (β2-β3, β5-β6, β10-β11, β13-β14,) located at the upper cap domain moved in the lefthanded direction ***(Figure 6D, E)***. These conformational movements led to a partial geometrical distortion that allowed protomeric bending in the pre-pore species ***(Figure 7A)***. However, such twisted or partly curved geometric distortion was more prominent in heptameric early pre-pore/oligomeric complexes ***(Figure 7A)***. These distorted oligomeric species also remained in predominant fractions (79%) isolated from pre-cooled RBC ***(Figure S14B)***. Unlike pre-pore states, the distorted heptameric states did not show any clear non-TMs β-barrel segments ***(Figure 7A)***. However, we were incapable of locating the density of the pre-stem segment inside the protomer associated with distorted states that predicted that the pre-stem detachment occurred before oligomer formation ***(Figure 6A)***. We hypothesized that either more cryo-EM data is required to resolve this pre-stem segment inside the protomer or due to inherent flexibility of pre-stem segment, we were unable to resolve this pre-stem segment.

**Figure 5:**
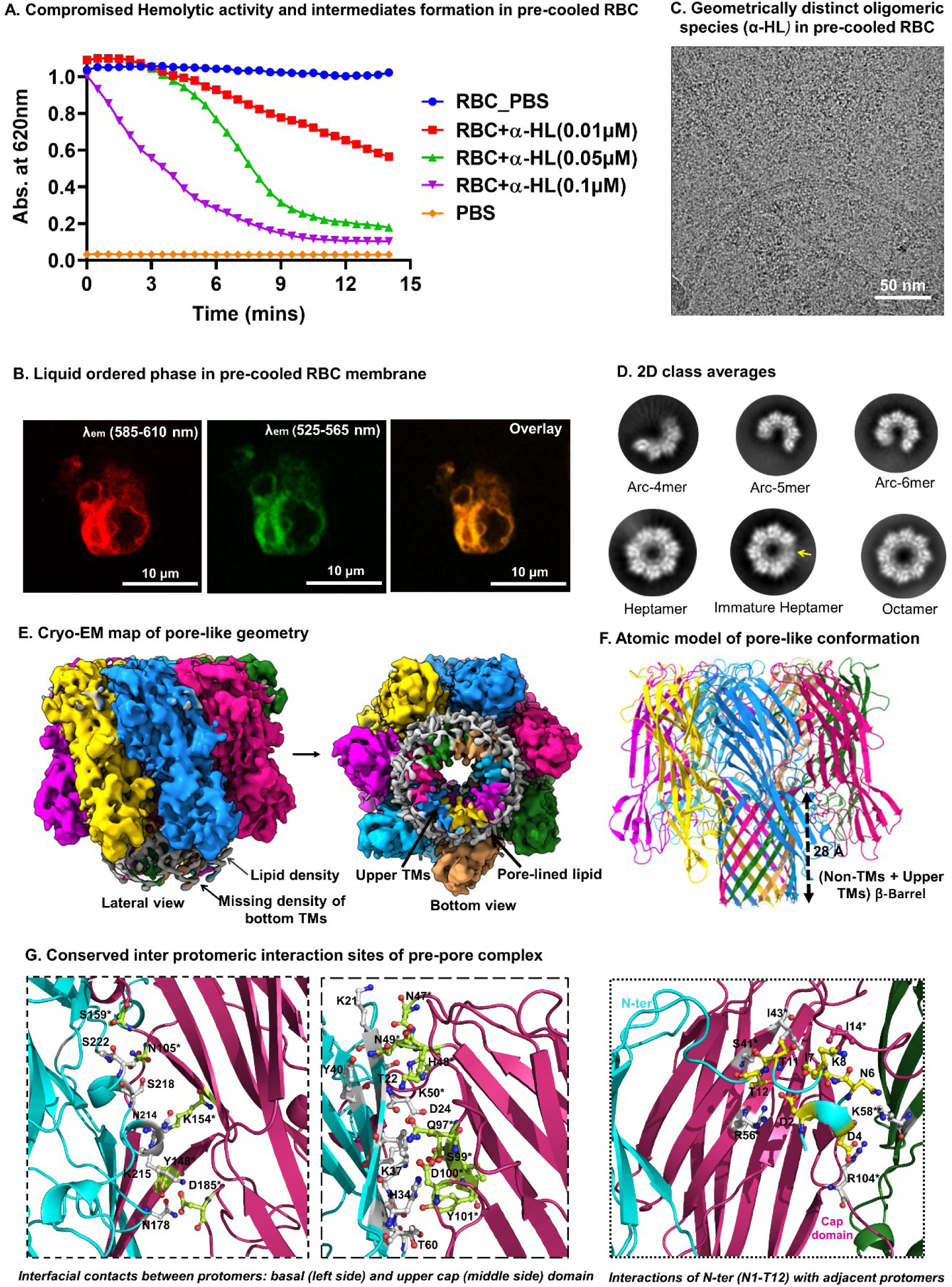
Lo phase of lipid bilayer restricted membrane perforation and prevented pore maturation of α-HL. **A.** A compromised haemolytic activity with pre-cooled RBC. **B.** Confocal images of pre-cooled RBC after Nile Red treatment showed a higher fluorescence intensity at 525-565 nm for a solvatochromic shift. **C.** The representative cryo-EM raw micrograph containing oligomeric particles originated from pre-cooled RBCs. **D.** Reference-free 2D class averages showed the existence of different geometrical species i.e., heptamer, octamer, and nonamer and arc-species. **E.** The cryo-EM 3D reconstruction of pore-like states (lateral and bottom view). Pore-lined lipid density (grey) coated the upper TMs channel and maintained the vestibular open channel. **F.** Atomic model of pore-like conformation having 2.8 nm long truncated β-barrel (missing density at lower TMs). **G.** Pore-like structure with conserved interfacial interactions like pore complex (dominated by polar residues; ball and stick) at upper and lower cap domains in between protomers, N-terminal segment of adjacent protomer maintaining several dipolar interactions with two vicinal protomers. The cartoon representations of adjacent protomers were shown in different colors (cyan, violet, and green respectively).

**Figure 6:**
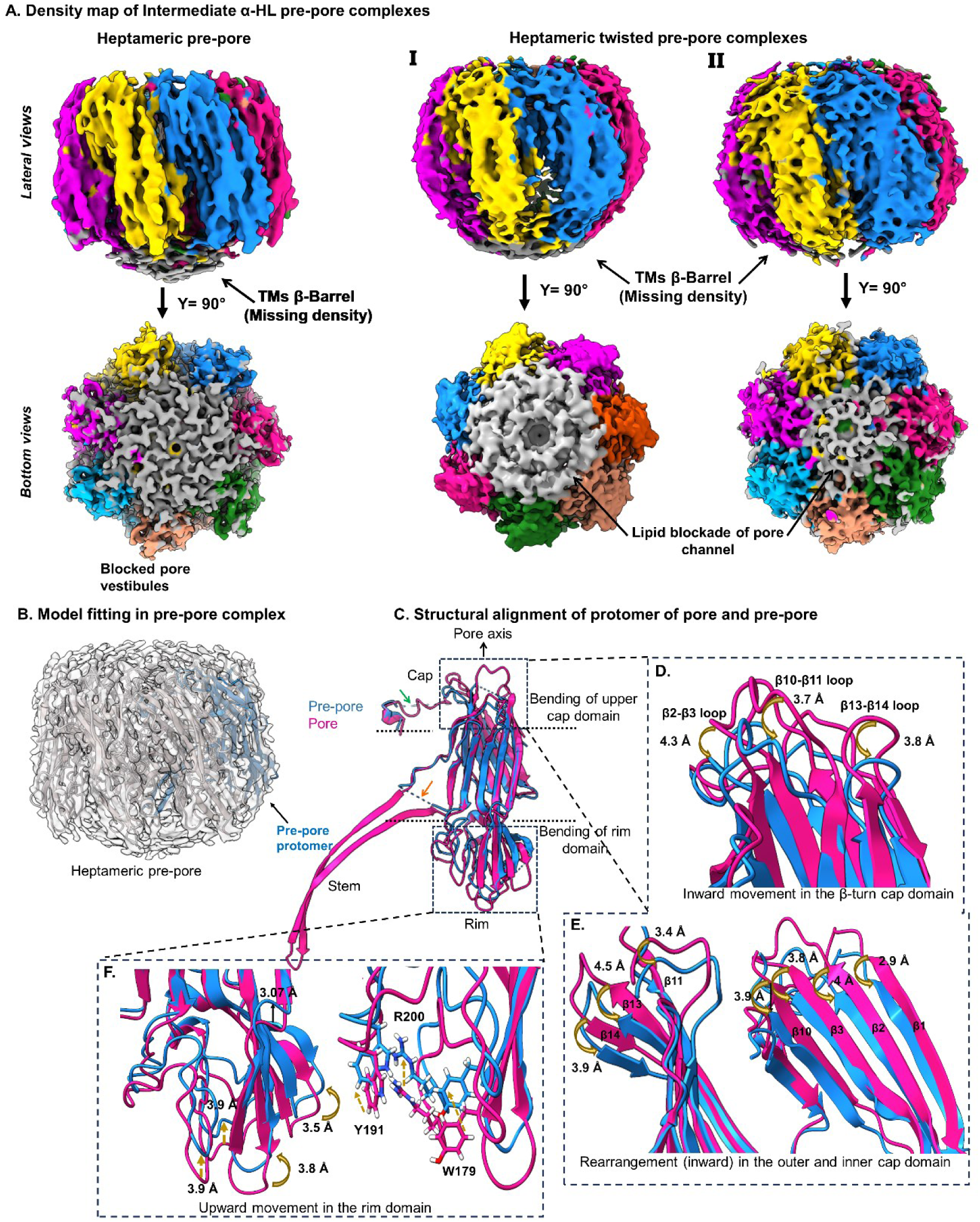
Conformational re-arrangements of α-HL protomers during pore formation and re-modulated lipid-protein interfaces in pre-pore complexes. **A.** Cryo-EM 3D map of heptameric pre-pore complex (left) and geometrically bend oligomeric intermediates (right; I, II). Lateral representation displayed discontinuous density of TMs segments, while the bottom views illustrated the blocked lipid densities (grey). **B.** Atomic model fitted with pre-pore EM map. **C.** The structural superposition of protomers from pore complex (violet) and pre-pore complex (blue). The cartoon representation suggested missing stem domain. Geometrical bending of upper cap and lower rim domains were highlighted (dotted box). **D.** Enlarge view represented a concerted inside and downward swing (w.r.to pore axis) of (β2-β3), (β10-β11) and (β13-β14) disordered loops in pre-pore complex. **E.** A similar trend of conformational shifts at top segments of inner (β1, β2, β3 and β10) and outer (β11, β13 and β14) cap β-sandwich domains. **F.** The cartoon representation of superimposed rim domains sighting a concerted inside and upwards movements of pre-pore structure. Lipid binding residues at lower rim domain highlighted in ball and stick model.

**Figure 7:**
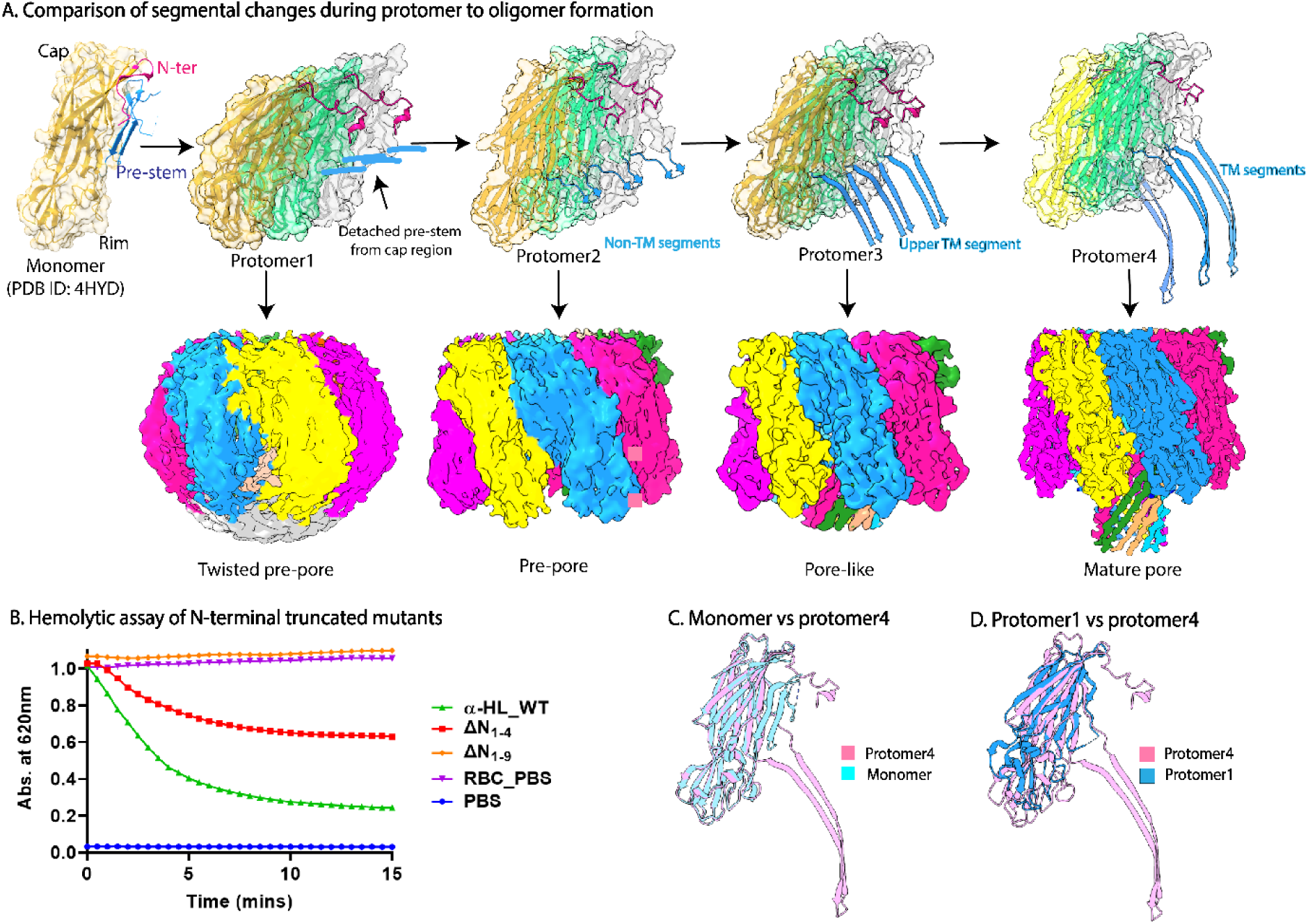
An overview of structural re-arrangements of water-soluble α-HL monomer to heptameric pore maturation and impact of N-terminal on pre-stem unfolding. **A.** The structural evolution from α-HL monomer to pore-forming protomers. Role of N-terminal segments and a plausible steric clash for pre-stem unfolding. Stepwise formation of upper TMs in pore-like structure and further conversion into a full TMs pore complex. **B.** A compromised hemolytic activity for partial (A1-D4) N-terminal truncated α-HL and completely abolished hemolytic (A1-T9) N-terminal truncated α-HL. **C.** The cartoon representation of structurally aligned monomer and α-HL protomer (pore) with a significant conformational change dominated in the pre-stem and stem domain. **D.** Geometrical deviation (inwards bending w.r.to pore axis) of protomer of a twisted pre-pore complex from the protomer of the mature complex.

The structural comparison of monomer and protomer in the pore complex of α-HL did not predict any large conformational rearrangement except pre-stem to-stem transition ***(Figure 7C)***. Surprisingly, we observed that the protomers embedded in pre-pore and early twisted oligomeric states maintained several other conformational movements of the monomeric toxin before oligomer formation ***(Figure 7A, D)***. An inward (with respect to the pore axis) bending of the upper cap domain and rim domain of protomers led to the formation of conformationally twisted early oligomeric and pre-pore geometry ***(Figure 7D)***. This was the first experimental evidence of large-scale conformational rearrangements identified for any small PFTs. Moreover, an increase in the orderedness of the lipid results in the prevention of barrel insertion, which generates large numbers of pre-pore species and other early oligomers. This further tells us about the findings of different intermediate states in the solution state.

## Discussion

In this current study, we aimed to understand the impacts of α-HL on innate immune cell and erythrocyte membranes. We showed for the first time that α-HL promotes necrosis in HL-60 cells with an increase in toxin concentration. Furthermore, we determined cryo-EM structure of heptameric α-HL toxin at atomic resolution form in the presence of rabbit erythrocyte, which was one of the first attempts to characterize small β-PFTs in the cellular environment. Despite having several crystals and cryo-EM structures of different^26,29–31^ pore-forming toxins, none of the structure of PFT has been characterized in real cellular environment. So far, our study highlighted several near atomic-resolution cryo-EM structures of intermediate pre-pore and pore-like structures in presences of invariable RBC membrane, which allowed us to understand pore formation mechanism by α-HL.

Previous studies showed that α-HL mediated hemolysis and cell membrane damage of erythrocyte and monocyte cells^10,11,32,33^. It was reported that α-HL activates necrosis and apoptosis cell death pathways in Jurkat and Lymphocytes cells^33,34^. Herein, we were interested to understand the toxin mediated effect on most abundant leukocyte and first line of host immune defense. Therefore, we demonstrated that α-HL initiated membrane damage and necrosis in HL-60 cells with an increase in toxin concentration. Herein, we observed that the self-association of α-HL on the host cellular plasma membrane eventually generated a membrane-inserted functionally active oligomer leading to the cell death process in cells. These phenomena were also observed with many other PFTs; like PFTs lysenin from *Eisenia fetida* specifically bound to U937 cells, aerolysin from *Aeromonas hydrophila* specifically interacted with U937 or THP-1 cells^35^, LLO and aerolysin both interacts with human intestinal epithelial cell line^36^. However, none of the studies described the detailed mechanism like our current study, where we demonstrated the detailed interactions of α-HL with innate immune cells and RBCs. Further, α-HL facilitated membrane protrusion from toxin bound area in cells and egg-PC/Chol LUVs that might help to reduce toxin load from the lipid bilayer. This was previously supported by several other studies; like Romero et al. showed that at the early stages of toxin invasions, the host cells could facilitate the defensive encounter process via removal of toxins embedded in the PC-enriched lipid microdomain from the plasma membrane^37^. Also, Gonzalez et al. showed that LLO and aerolysin both interacts with human intestinal epithelial cell line, although membrane recovery rates were different for both the toxin^36^.

There are few crystal structures of α-HL were characterized where none the studies were performed using lipid environment or cell membrane^29,30,37^. Recently pore and intermediate pre-pore conformations of *Vibrio Cholerae* Cytolysin (VCC) on phospholipid membrane were also characterized in the presence of lipid environment^31^.

Thus, in this current study we determined the cryo-EM structure of α-HL form in the presence of rabbit erythrocyte, which was one of the first attempts to characterize any PFTs in the physiological cellular environment. There were several cryo-EM structures that were resolved by cryo-electron tomography (Cryo-ET) of cells with large oligomeric protein complexes, spike proteins of viruses or virus particles attached with cell^38–40^. But none the studies focused on small pore forming toxins attached with cells. In these circumstances, our approaches were distinctive from others, which might be employed to characterize many other PFTs and effector proteins in cellular environment.

In order to understand the lipid-toxin crosstalk in a vivid manner, we characterized the cyro-EM based structural analysis of α-HL in the different chemical and physical states of phosphatidylcholine lipid membrane. In our study, we resolved heptameric pore and pre-pore species in the presence of 10:0 PC as well as a rapid lysis of the 10:0 PC vesicle was observed upon toxin treatment. This could be due to the relatively small bilayer thickness (~2.6 nm) of the 10:0 PC LUVs suffered by bilayer stretching while stabilizing the hydrophobic part of ~3.6 nm TMs of mature heptameric pore species. In our previous study, we observed membrane rupture in case of HlgAB from *Staphylococcal aureus,* which formed octahedral shaped pre-pore super-assemblies^41^. Similar, higher order assemblies we could also observe along with circular oligomer in 10:0 PC condition, which could arise from the individual pre-pore species. Cryo-EM structure of the heptameric pre-pore complexes originated from 10:0 PC contained thick lipid densities located at vestibular fenestration of blocked channel, which is correlating the previously reported pre-pore state obtained in VCC^31^.

In addition, we provided a detailed glimpse of how the membrane fluidity could significantly re-model the structural metamorphosis of α-HL. The incorporation of planner cylindrical-shaped lipid SM introduced Lo-phase behavior inside bio-membranes^14,42^. Stoddart et al., showed that truncation of β-barrel region up to 27Å did not diminish transmembrane activity of the oligomeric complex^43^. Thus, the formation of pore-like states in 10:0 PC/SM LUVs maintained the functional activity like mature pore complex. Furthermore, previous reports had predicted the membrane fluidity played a crucial role in membrane perforation and stabilization of transmembrane barrel of different toxins^3^. Interestingly, the pre-cooled egg-PC/Chol and pre-cooled RBC decreased the pore-induced leakage and membrane lysis due to a significant increase in Lo-phases in their respective model and plasma membranes. Due to limited and restricted membrane perforation, we observed several intermediates oligomeric states of α-HL. The different geometrical and conformational states of the toxin confirmed a sequential stepwise oligomerization pathway. The formation of an intermediate pore complex of α-HL might be generated due to the dynamic process of protomer association, the relative occurrence of any such intermediate pre-pore complex and early oligomeric states solely depends on the physiochemical states of lipid bilayer. Therefore, to combat host battle while exposed to an external endeavor like PFTs, the SM/saturated enriched lipid microdomains (Lo) of the host plasma membrane could prevent pore formation while large fluidic PC lipid membrane components (Ld) could facilitate the membrane protrusion to get rid of excess toxin load.

In summary, our current study elucidates that α-HL not only lysis the rabbit erythrocyte, but it is also capable of damaging cultured HL60 cell and RAW264.7 plasma membrane. We successfully resolved several intermediate and complete pore structures of α-HL at the atomic resolution in the presence of rabbit erythrocyte and pre-cooled membrane stabilized RBCs. This is the first report to capture all the intermediates states of α-HL in real cellular environment. Therefore, our study provides the first direct visualization of oligomeric assembly and membrane damaging process by α-HL.

## Materials and methods

### Cloning, overexpression, and purification of α-HL

α-hemolysin (α-HL) codon-optimized gene (Genscript, USA) was cloned in a C-terminal 6x His-tagged pET26b(+) vector. The synthesized α-HL gene was transformed and overexpressed into *E. coli* BL21 (DE3) strain cells (Novagen) using Luria Broth (HiMedia) growth media to an OD_600_ of 0.6-0.8 at 37°C. Isopropyl-b-D-thiogalactoside (IPTG, HIMEDIA) at 0.5 mM concentration was added into the cultured cells at 25°C for 12 hours to overexpress the protein. The cultured cells were harvested at 4500 rpm for 20 mins and the pellet containing cells was further dissolved in lysis buffer (50 mM Tris-HCl buffer pH 8, 150 mM NaCl, 2% glycerol), 7mM imidazole, 2 mM phenylmethyl sulphonyl fluoride (PMSF), 1mM DTT and 0.8mg/ml lysozyme. After 30 minutes incubation after adding lysis buffer, cells were further lysed using 30 min sonication (20kHz, amplitude 25%, Fisher Scientific) and followed by centrifugation for 40 minutes at 13000 rpm in order to remove insoluble cellular debris.

The supernatant solution was incubated with pre-equilibrated with lysis buffer containing 7mM imidazole Ni–NTA agarose beads (QIAGEN) and kept for binding for 3 hours on a rotating platform at 4°C. After passing the supernatant flow through in a chromatographic column, the beads were washed with a wash buffer containing 50 mM Tris-HCl buffer pH 8, 150 mM NaCl, 2% glycerol, and 20-60 mM imidazole (Sigma Aldrich) gradient concentration. Finally, the α-HL protein was eluted at a 300 mM imidazole concentration in the lysis buffer.

The eluted protein using Ni-NTA chromatography was buffer-exchanged before size exclusion chromatography. To isolate monomeric toxin the protein was loaded into the 10/300 GL column (Cytiva) packed with Superdex 200 beads pre-equilibrated with buffer compositions of 50 mM Tris-HCl buffer pH 8, 150 mM NaCl. The isolated protein was concentrated using Amicon® Ultra centrifugal filter with a molecular cutoff 10 kDa (Amicon Ultra-4 Centrifugal filter unit 10K, Merck). The SEC peak fraction for recombinant α-HL was further loaded into 10% SDS-PAGE (Coomassie Brilliant blue R 250, Merck) to confirm its purity. Precision Plus Protein™ Dual Color Standards (Bio-Rad) were used as a protein molecular weight marker in SDS-PAGE.

### N-terminal truncated clones generation of wild type α-HL

For the N-terminal deleted truncation (A1-D4 and A1-T9 amino acids respectively) construct generation, PCR was performed with an initial thermal denaturation at 98°C for 30 s, a follow-up primer annealing at 58-65°C for 30 s, DNA synthesis at 72°C for 5 min, and a final extension at 72°C for 10 min. The PCR products were ligated using ligation mixture. The ligated DNA mixture was purified and transformed in *E. coli* DH5α cellsThe generated N-terminal truncated clones were further confirmed by DNA sequencing. The N-terminal truncated clones of α-HL were overexpressed and purified using the same method described for wild type α-HL.

### Haemolytic assay

About 1 ml of whole blood was isolated from *Oryctolagus cuniculus* and was centrifuged several times at 1000 rpm for 10 minutes using 1X Sodium Phosphate buffer saline (PBS) containing 137 mM Sodium Chloride (Qualigens), 1.8 mM Potassium Di-hydrogen Orthophosphate (Qualigens), 10 mM Disodium Hydrogen Orthophosphate (SD Fine Chemicals), 2.7 mM Potassium Chloride (Sigma Aldrich) to isolate plasma components free intact RBCs for haemolysis assay with α-HL and N-terminal truncated proteins. 1X PBS buffer solution was used to prepare 10% RBCs solution. 1% Triton-100X detergent solution was used as a positive control for the complete haemolysis process while 1X PBS buffer was used as a negative control for the experiment. RBCs were treated with different (μM) concentrations of α-HL and incubated in constant shaking conditions (160 rpm) at 37°C for a duration of 15 minutes. The optical density (O.D) of the RBC cells in different conditions of toxin treatments was measured at 620nm using a Varioskan Flash microplate reader (Thermo Scientific). Haemolytic assay of the N-terminal truncated proteins were also performed in similar manner.

### Cell death analysis using flow cytometry

HL-60 leukemia cells were maintained in RPMI 1640 media supplemented with 2mM Glutamine and 10% Fetal Bovine Serum in 37°C and 5% CO_2_ incubator. Approximately 1*10^6^HL-60 cells were taken and were washed twice with the wash buffer containing PBS with 3% BSA (pH 7.4). The cells were then treated with 10 nM, 100 nM, and 1 µm of α-HL for 1 hour 30 minutes at room temperature. Following treatment with α-HL, the cells were washed again twice with the wash buffer and were incubated for 30 minutes after subjecting to single and dual labelling with Annexin V-FITC or Ethidium homodimer III (EthD-III) at 4^0^C under constant shaking condition. After labelling, the cells were again washed twice with wash buffer and were analyzed by Flow Cytometry using BD Aria III instrument. For analyzing the cells by flow cytometry, 530/30nm band pass filter was used for Annexin V-FITC and 585/42nm band pass filter was used for Ethidium homodimer III (EthD-III).

### Confocal imaging

To analyze the cytotoxic impacts in cultured HL-60 cells and RAW264.7 cells were incubated with toxin at a 50 nM concentration of α-HL and were visualized under Zeiss ELYRA 7 allows ApoTome imaging using 40X oil objective with Lattice SIM super-resolution microscope in the presence of lipophilic membrane dye Nile Red and DNA staining dye DAPI. An excitation laser source at 405 nm and 488 nm was used for DAPI and Nile Red fluorescence imaging respectively. The emission band pass filter for DAPI and Nile Red was 480-510 nm and 570 nm respectively. The 3DSIMsettings using ZEN software (ZEISS India) corresponded to a periodic grid of period 23mm for 488nm and 28mm for 555nm, 3 rotations, and 5 translations. Leica Falcon SP8 was also used to perform all the confocal data for fluorescently labeled liposomes and cell membranes incubated with toxins. Nile Red provided a solvatochromic diverse emission spectrum after binding on Lo/Ld phage of lipid and RBC plasma membrane^45^. Nile Red was added in the media of cultured cells (35mm dish) and imaged immediately. Followed by addition of toxin and DAPI simultaneously. The captured z slices were analyzed to monitor toxin induced damage of cell membrane and nucleus. The images were analyzed using Fiji ImageJ^46^.

### Preparation of lipid vesicles

Phosphatidylcholine (10:0) or 10PC, 14:0 PC, 18:1PC), egg-PC/ Cholesterol (3:1 molar ratio), egg-PC/ Sphingomyelin (3:1 molar ratio) composite lipids were taken for the preparation of liposome separately by lipid extrusion method. The lipids were first dissolved into 1 ml chloroform and further allowed for evaporation of the residual chloroform from respective lipids. The layer of individual lipid mixture was further dissolved gently into 300μl total volume of 1X PBS buffer (pH 7.4) and further incubated for 30 minutes at an above transition temperature (T_m_), 50°C. The lipid solutions were further extruded through 1μm and 200 nm pore-size polycarbonate membranes (PC Membranes 0.2μm, Avanti Polar Lipids) 15 times to form Giant unilamellar vesicles (GUVs) and large unilamellar vesicles (LUVs). The final extruded solution containing 4 mM LUVs and GUVs were used in further experiments. Avanti Polar mini-extruder apparatus contained 10mm Filter supports, and Hampton syringe for liposome preparation.

### Total internal reflection fluorescence microscopy (TIRFM)

For fluorescence imaging of liposomes under TIRF microscope, a glass slide and coverslip passivated with polyethylene glycol (PEG) was used. The preparation of such surfaces involves a series of steps described previously^47^. The flow channels used for imaging were prepared by sandwiching a double-sided tape between the PEG-passivated glass slide and coverslip. A small fraction (1:100) of the PEG molecules carry biotin for immobilization. The ends of the channels were sealed with epoxy glue and the holes drilled in the glass slide enabled buffer exchange. We then incubated the flow channels with 0.5 mg/mL streptavidin (Sisco Research Labotaries) in T50 buffer (50 mM Tris-HCl, 50 mM NaCl, pH 7.5) for 10 minutes. Then the channels were washed with 500 µL of T50 buffer before immobilizing the liposomes.

We flowed in 30 µL of 4ME16:0 NBD-PE or egg Liss Rhod-PE (Avanati polar lipid) labelled biotinylated liposomes with encapsulated Rhodamine B dye at a flow rate of 10 µL/min. During the flow, the biotinylated liposomes get tethered onto the surface via biotin-streptavidin interactions. A 50 µL of PBS (pH 7.4) was then flowed in at the same flow rate to wash out the unbound liposomes.

The image acquisition was set up to image both the bound liposome (488 nm channel) as well as the rhodamine B dye inside the liposome (561 nm channel). Finally, we flowed in 10 nM of the toxin α-HL and recorded the changes fluorescence intensities in both the channels. The images were analyzed using Fiji ImageJ^46^.

### Fluorescent labelling of wild type α-HL

S278C, a single cysteine mutant of α-HL was used for labeling Atto 647N maleimide fluorescent dye (Sigma Aldrich). Initially, a 20 mM stock solution of the Atto 647N maleimide dye was prepared in DMSO immediately prior to use. 50 µM mutant protein solution was mixed with excess amount of Atto dye in respective binding buffer (Sigma Aldrich) and incubated overnight at 4°C in a dark place. The excess free unbound dye was removed by 10 kDa molecular filter (Amicon Ultra Centrifugal filter) prior to further use.

### Negative staining sample preparation and visualization by TEM

Three microliters sample of monomeric toxin incubated with different liposome {Phosphatidylcholine (10:0) or 10PC, 14:0 PC, 18:1PC), egg-PC/ Cholesterol (3:1 molar ratio), egg-PC/ Sphingomyelin (3:1 molar ratio)} solutions were applied to a thin film of carbon-coated copper grids (Carbon film 300 mesh, Copper, Electron Microscopy Sciences) at room temperature, which was glow discharged for 30 seconds, 20 mA freshly. These samples were allowed to settle down on grid for 1 minute, before removal of excess buffer. 1% uranyl acetate (Polysciences, Inc) was used for staining, and followed by blotting to remove excess stain from the grid. These negatively stained samples on carbon-coated copper grids were then visualized under a 120 kV Talos L120C transmission electron microscope equipped with a bottom-mounted Ceta camera (4000 x 4000 pixels) at magnification ranging between x57000-x92000.

### Data Processing for NS-TEM micrographs

Different particle projections were manually picked from 2D raw NS-TEM micrographs using e2projectmanager.py (EMAN2.91^48^) followed by the extraction of particles with a 160 pixel box size and a processing 56 Å mask diameter from raw micrographs using e2boxer.py. An average of 6000-7000 particles from each dataset were further classified into 2D-class averages using reference-free 2D classifications in EMAN2.91^48^, RELION2.0^49^, and SIMPLE^50^ programs.

### Cryo-EM Sample Preparation

R1.2/1.3 300 mesh copper grids (Quantifoil; Electron Microscopy Sciences) were glow discharged for 90 seconds at 20 mA before cryo-sample freezing process. For the RBC-α-HL complex, α-HL (1 mg/ml) was mixed with rabbit erythrocytes. After incubation, we centrifuged the mixture at 13,000 rpm for 30 minutes. Three microliters of supernatant solution from that cell-protein mixture were applied onto the freshly glow discharged grids and immediately blotted for 8 secs with blot force of positive 25 using FEI Vitrobot Mark IV plunger. The grid was plunged into the liquid ethane instantly after blotting. For pre-cold RBC, the RBC was incubated with the α-HL in ice and then underwent the same freezing procedure for cryo-EM data collection. The toxin-treated different composites liposomes were also cryo-frozen in a similar freezing parameter.

### Cryo-EM Data Collection

Cryo-EM data were acquired using 200 kV Talos Arctica transmission electron microscope (Thermo Scientific™) equipped with K2 Summit Direct Electron Detector (Gatan Inc). Movies were recorded automatically using Latitude-S^51^ (Digital Micrograph - GMS 3.5) at nominal magnification of 54,000x at the effective pixel size of 0.92 Å. Micrographs are collected in counting mode with a total dose of 50 e^−^/Å^2^, with an exposure time of 8 sec distributed for 20 frames. A total of 643, 1427, 1170, 681, and 1521 micrographs were acquired for the RBC_α-HL, egg-PC LUVs_α-HL, 10:0 PC LUVs_α-HL, 10:0 PC/SM LUVs_α-HL, and RBC_cold_α-HL protein complexes respectively.

### Cryo-EM Data Processing

Single-Particle Analysis (SPA) were performed for the acquired cryo-EM data using the RELION version 3.1^52^. In the pre-processing step, the drift and gain corrections of the individual movies were performed with MotionCor2^53^. The micrographs were chosen for an estimation of Contrast transfer function (CTF) parameters using CTFFIND 4.1.13^54^. Initially, particles were manually picked using RELION 3.1^52^ for RBC_ α-HL dataset followed by 2D classification. The best 2D class averages were selected as template for auto-picking for all the collected datasets. The particles were extracted with the box sizes of 280-pixel, calibrated pixel size of 0.92 Å for all the membrane-bound toxin complexes. Several rounds of 2D classification were run each time to remove junk particles. The best 2D classes were selected for 3D classification with C7 symmetry ***(Figure S9, S11B, S13)***. The ab-initio model generated from one good class of a 2D class averages and the converted map from the crystal structure (PDB ID: 7AHL^29^) was filtered to 40 Å to use as a reference for 3D classification. The 3D class averages of various heptameric conformations were subjected to movie refinement, which includes the estimation of beam tilt, anisotropic magnification, and per-particle CTF refinement for defocus and astigmatism. The sharpening for the 3D auto-refined maps was performed with RELION 3.1 and PHENIX^55^. Global resolution of Fourier Shell Correlation (FSC) was estimated at the threshold of 0.143^56^ and the estimation of a local resolution was performed with ResMap^57^, using auto-refined half maps.

### Model building and structure refinement

Cryo-EM maps were initially docked with PDB ID 7AHL^29^ and 4YHD^58^ models in UCSF chimera to use as a template for atomic model generation for heptameric pore, pore-like, pre-pore complex and twisted heptameric pre-pore complex respectively. Automated model building and further refinements were iteratively performed with Phenix Real Space Refinement in using coot^59^ and Phenix^60^. The phenix.real_space_refine models and part of the phosphatidylcholine were fitted to respective cryo-EM density maps using UCSF ChimeraX^61^.

### Analysis and Visualization

The structural statistics for Cryo-EM map and atomic model were analyzed and validated using Phenix, UCSF ChimeraX and EMringer^62^, Molprobity^63^, respectively. Surface coloring based on electrostatic surface potential and hydrophobicity scale were performed using UCSF ChimeraX^61^. The structural comparison and model alignment was performed after overlapping the protomers using UCSF ChimeraX “MatchMaker” tool. The detailed structural analysis were performed using UCSF Chimera, ChimeraX^61^ and PyMOL.

## Acknowledgments

We acknowledge Department of Biotechnology (DBT), Department of Science and Technology (DST), Council of Scientific and Industrial Research (CSIR), and Ministry of Education (MoE), India for funding and cryo-EM facility at IISc, Bengaluru. We acknowledge DBT-BUILDER Program (BT/INF/22/SP22844/2017) and DST-FIST (SR/FST/LSII-039/2015) for National Cryo-EM facility at IISc, Bengaluru. A.C. acknowledges Council of Scientific and Industrial Research (CSIR) Grant Number-09/0079(13652)/2022-EMR-I) for doctoral fellowship. A.R. acknowledges the financial support from the Ministry of Education (MoE) (Grant Number-STARS-1/171). We acknowledge Science and Engineering Research Board, India (Grant No. CRG/2022/002674) for financial support. SD thanks STR/2022/000006 SERB-STAR award for financial support. We are grateful to Ms. Aparna Asok for her help in performing flow cytometry-based experiments. We thank Mr. Prathibhan KV, and Mr. Gourab Chatterjee for help in TEM imaging and protein purification respectively.

## Author contributions

A.C., A.R. and S. D. designed the experiments; A.C., A.R., P.P.D., B.M, generated mutant, truncated clones and purified proteins; A.C., D.C., A.R., B.G. performed cell-based experiments; A.C., A.R. performed biophysical characterizations; A.C., T.S., M.G., S.D. performed TIRF based imaging and image analysis; A.C., S.D. performed TEM based experiments; A.C., P.K., S.D. performed cryo-EM sample preparation and data acquisition; A.C., A.R., S.D. performed cryo-EM data processing and analysis; A.C., A.R., D.C., M.G., S.J. S.D. wrote and reviewed the manuscript.

## Ethics approval

Rabbit plasma was provided by Institutional Animal Ethics Committee (IAEC), Indian Institute of Science, Bangalore (CAF/Ethics/881/2022).

---

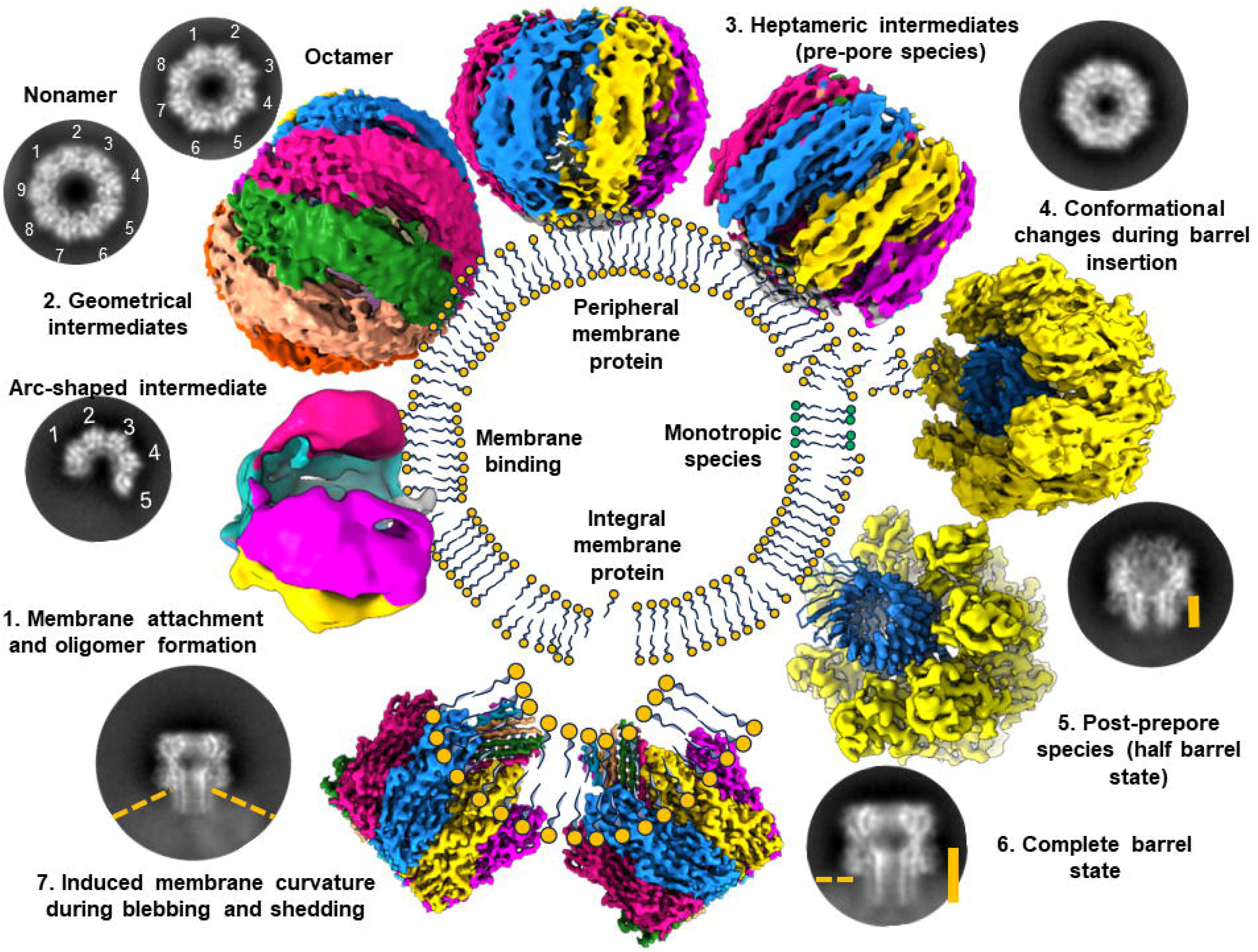

