## Supplemental figures for "Crosstalk between plasma membrane and *Staphylococcus* α-hemolysin during oligomerization"

[$]Equally contributed.

### Supplemental Figures and Legends

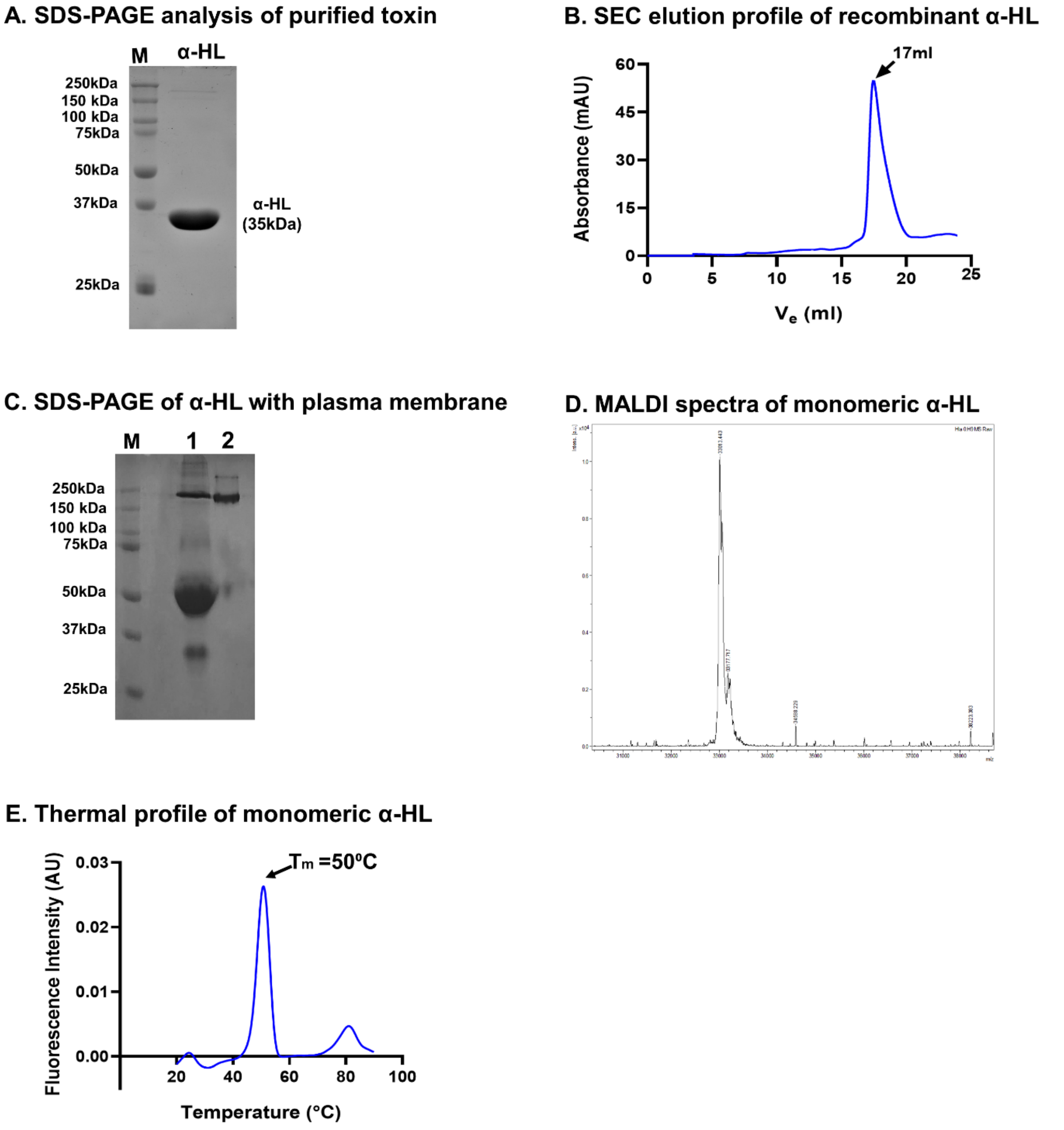


***Figure S1: Preliminary purification and characterization of α-HL.***

***A.*** *SDS-PAGE analysis of gel filtrated freshly purified recombinant α-HL monomer (M-Precision Plus Dual Color Protein Standard).* ***B.*** *Size exclusion chromatography (Superdex$^{TM}$ S200 increase 10/300 GL) profile of recombinant α-HL monomer.* ***C****. SDS-PAGE profile of protein contents presents in the supernatant fraction of α-HL treated HL-60 cells (lane 1) and rabbit erythrocytes (lane 2) respectively.* ***D.*** *MALDI spectra analysis of recombinant α-HL showed a molecular weight of intact protein of ~35kDa.* ***E.*** *Thermal stabilization/denaturation spectral profile of the monomeric toxin.*

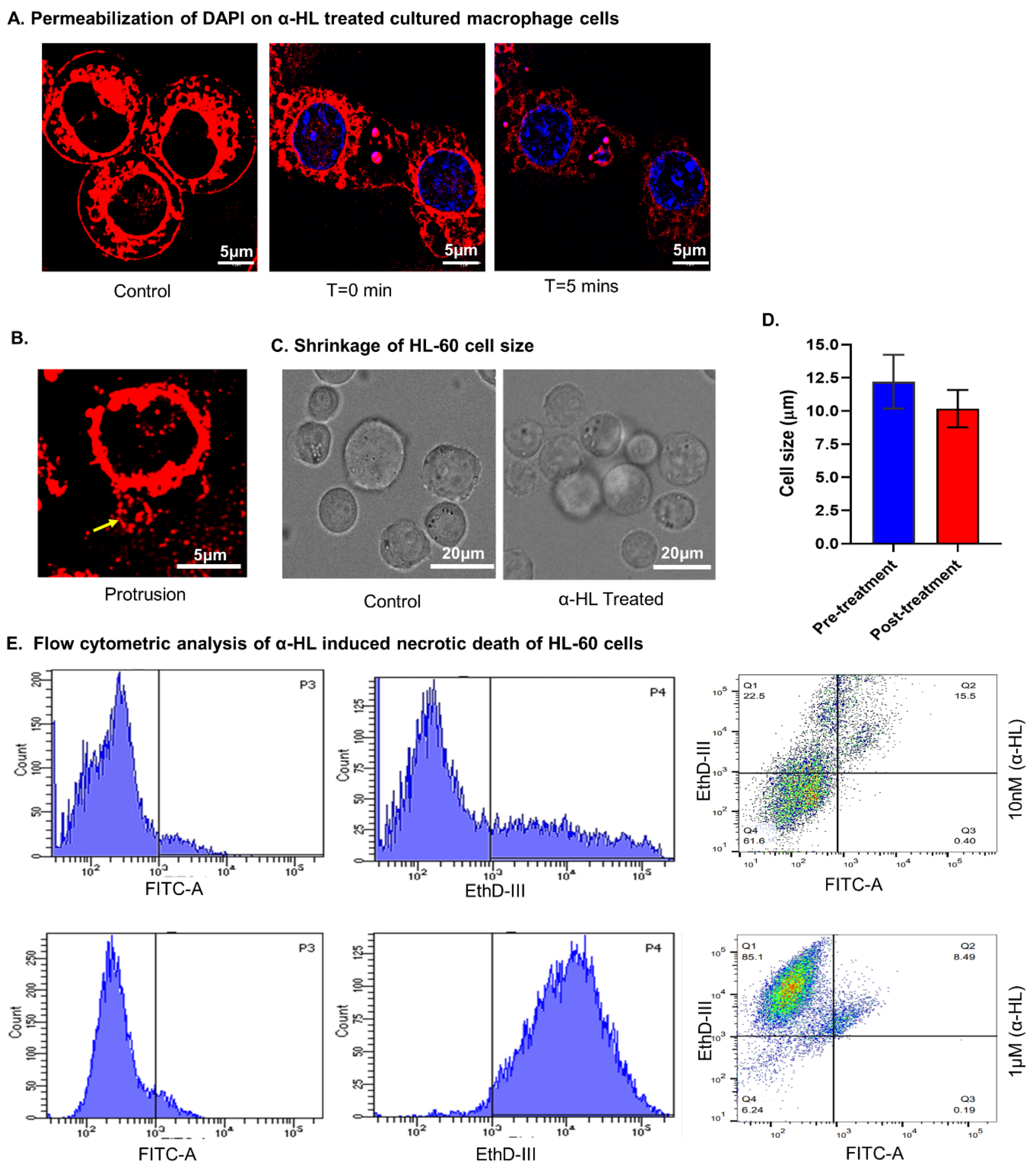


***Figure S2: Effect of α-HL on innate immune cells.***

***A.*** *Permeabilization of a DNA binding dye (DAPI) in cultured macrophage cells (RAW264.7) due to toxin treatment as control (left) and observed at different time intervals after 50nM toxin(α-HL) treatment. An overall increase in intracellular DAPI intensity was observed over 5 minutes.* ***B.*** *Cultured HL-60 cells were incubated with α-HL for 5mins. The protruded body was detected after toxin treatment (yellow arrow).* ***C.*** *Cell size shrinkage along with membrane protrusion at 50mM toxin concentration and clustering of cells at high toxin concentration (500nM) were also observed.* ***D.*** *The relative change in cultured HL-60 cell size after treatment of α-HL toxin.* ***E****. FACS analysis of HL-60 cells treated with α-HL. Cells treated with sub-nanomolar to micromolar concentrations of toxin showed a significant increase in cellular necrosis.*

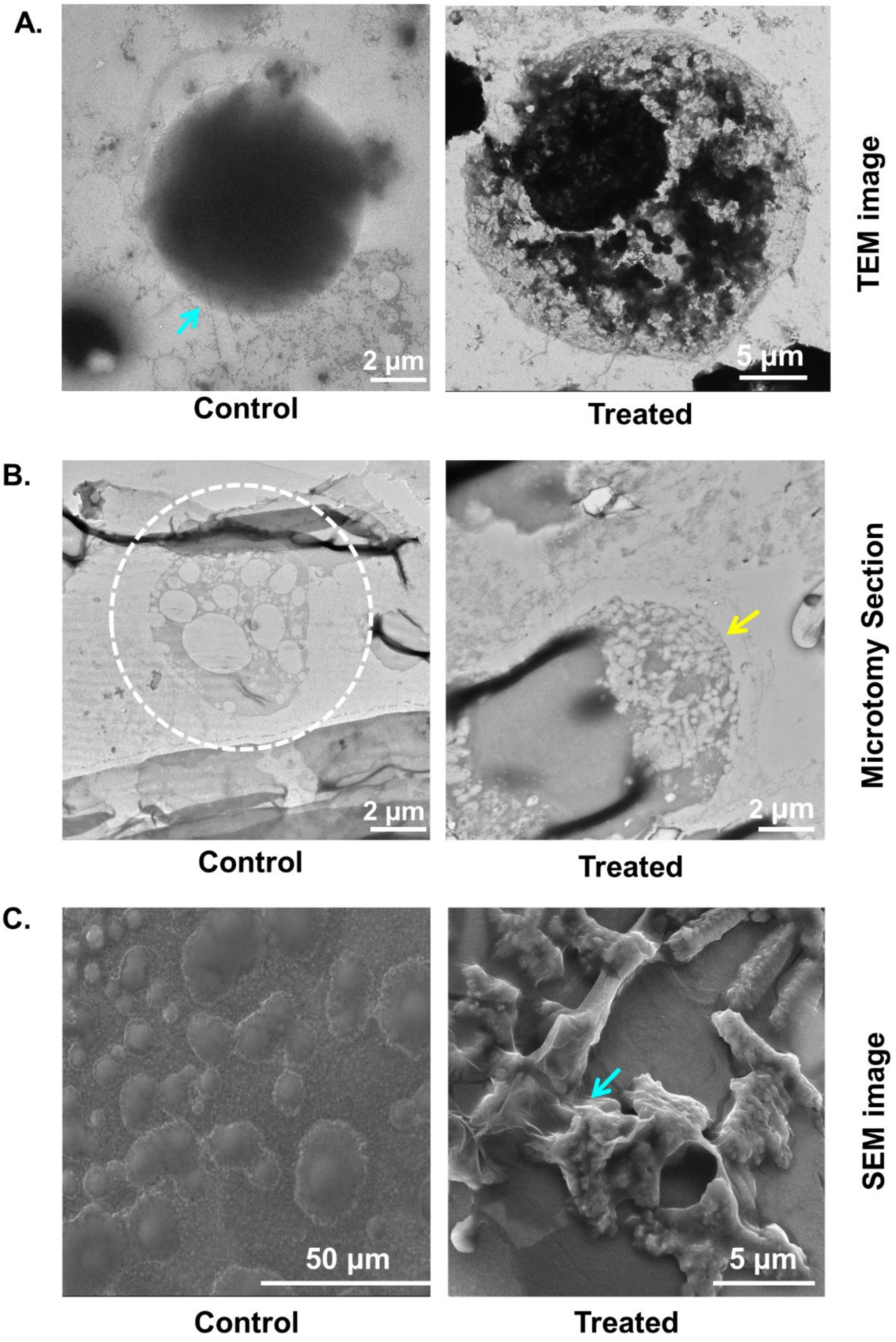


***Figure S3: α-HL induced morphological rearrangements of HL-60 cells.***

***A.*** *Transmission Electron Microscopic (NS-TEM) image showed spherical morphology of HL-60 cell (control; cyan arrow) and α-HL (50nM) treated partially damaged HL-60 cells.* ***B.*** *Ultra microtomic electron micrograph of granular HL-60 (white encircle) cell and partly lysed state (yellow arrow) after α-HL incubation.* ***C.*** *Scanning Electron microscopic (SEM) showed a distinct morphological change of the cells (cyan arrow) after incubation with 50nM monomeric toxin.*

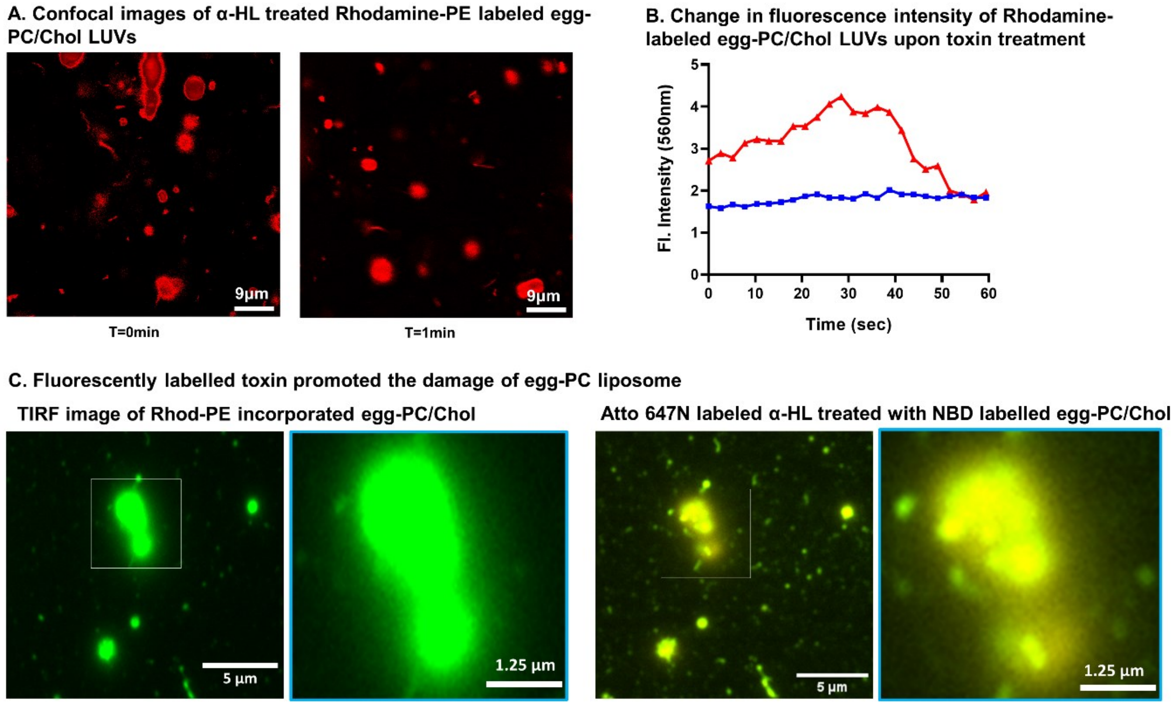


***Figure S4: Toxin-mediated partial lysis of egg-PC liposome.***

***A.*** *Confocal images of control Rhod-PE ($\lambda_{em}$= 570 nm) loaded egg-PC/Chol LUVs/GUVs and α-HL treatment (1 min).* ***B.*** *A relative change in fluorescence intensities at different intervals of toxin-treated (red) and control vesicles (blue) is shown respectively.* ***C****. TIRF image showing a pseudo green fluorescence of Rhod-PE tagged egg-PC/Chol LUVs/GUVs (left). Binding and co-localization of atto488 fluorescent-tagged 278C α-HL ($\lambda_{em}$= 540 nm) on Rhod-PE tagged egg-PC/Chol liposome represented in an enlarged view (right).*

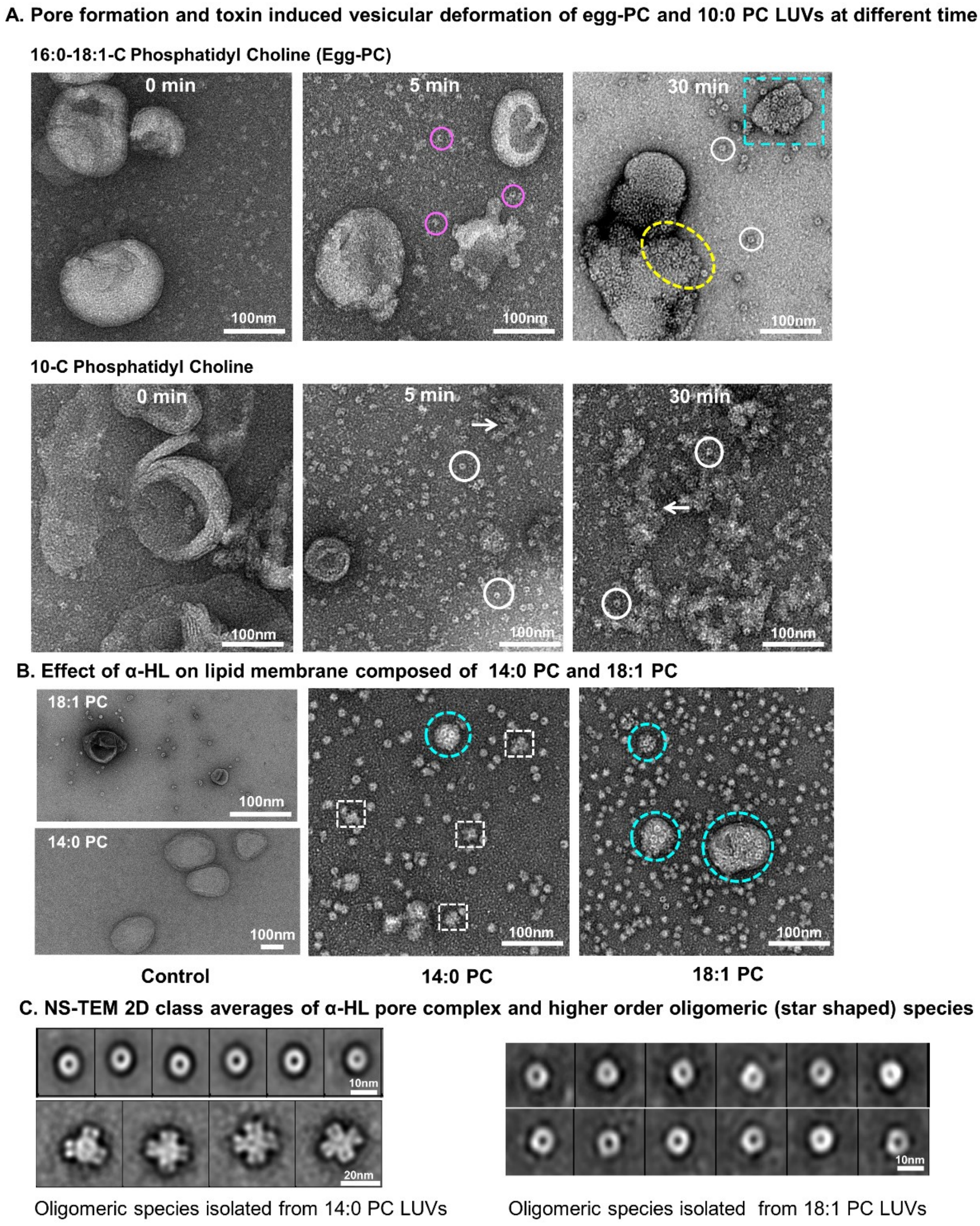


***Figure S5: Different modes of toxin (α-HL) mediated lipid membrane re-modulations.***

***A**. NS-TEM micrographs showed time laps of pore formation followed by membrane protrusion on egg-PC/Chol vesicles. Monomeric α-HL (50nM) treated LUVs of egg-PC/Chol (0min) initiated the oligomer formation (Pink encircled) upon incubation for 5-minute intervals. Fragmentations of lipid vesicles in terms of protruded bodies loaded with PFTs (cyan box), localizations of pore species on LUVs (yellow encircle), and oligomeric species detached from vesicle surface (white encircles) after 30 mins post toxin treatment. Formation of α-HL pore complex and consecutive lysis of 10:0 PC vesicles after toxin incubation for 5mins and 30mins respectively. **B.** Formation of oligomeric pore complex covered on the small protruded 18:1 PC micro-vesicles (cyan encircle). NS-TEM imaging also showed a few higher-ordered oligomeric species (white dotted box) originated from 14:0 PC LUVs. **C.** The representative NS-TEM 2D class averages of oligomeric states of α-HL and higher ordered oligomeric species of α-HL from 18:1 PC and 14:0 PC respectively*

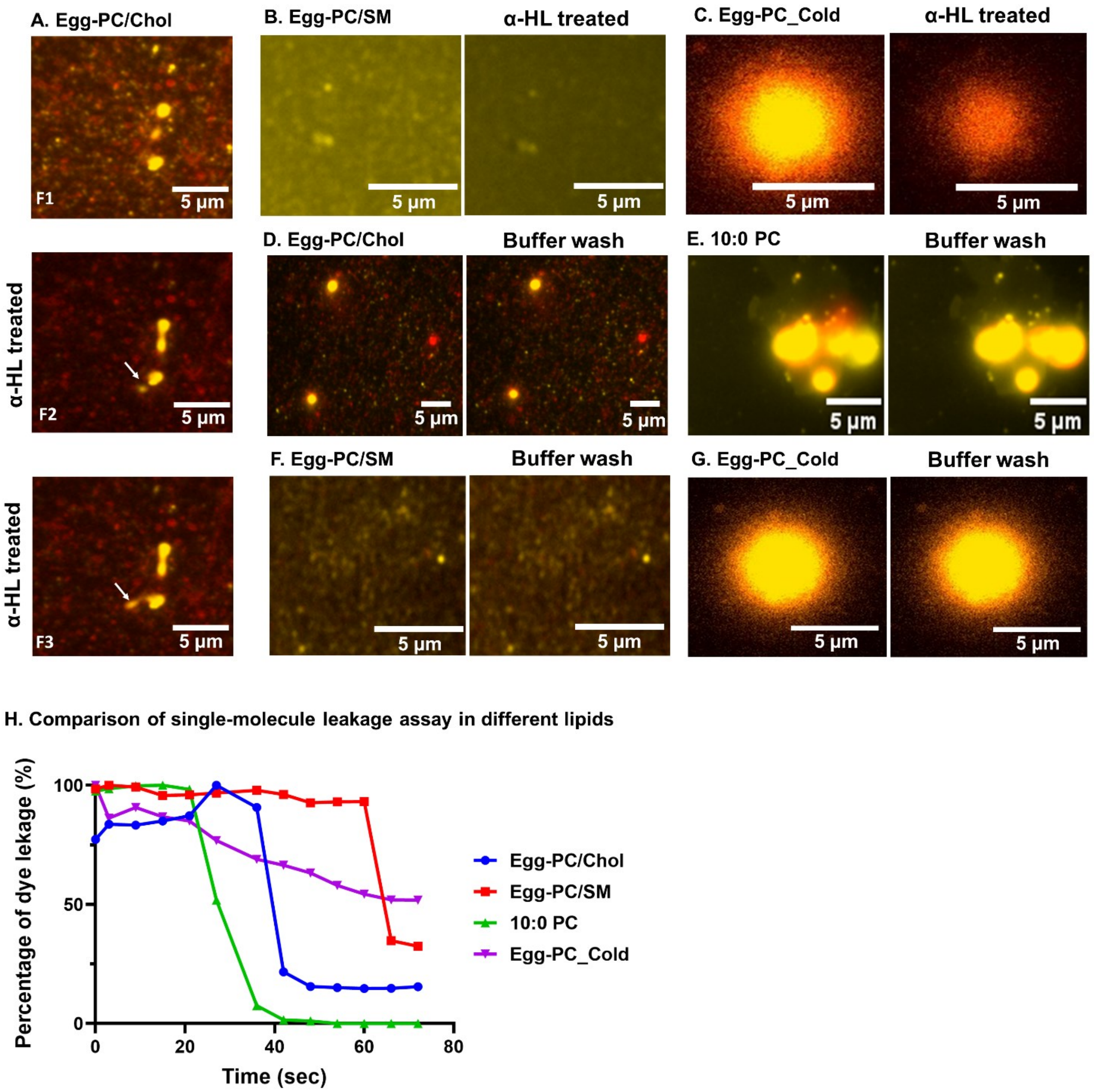


***Figure S6: α-HL induced rhodamine leakage assay with different composites vesicles.***

***A.** Protrusion of egg-PC/Chol lipid vesicle after α-HL treatment. **B-C.** Total Internal Reflection Fluorescence (TIRF) microscopic imaging of toxin-treated NBD-PE labeled eggPC/SM, and pre-cooled eggPC/Chol lipid vesicles shown in pseudo colour (red), filled with rhodamine (in yellow) showed a decrease in the fluorescence of Rhodamine dye. **D-G.** The relative fluorescence intensities for both fluorophores remained unchanged while washing with buffer through the flow-channel-bound liposomes. **H.** Comparison of single-molecule based different composites vesicle encapsulated rhodamine leakage assay showed significant delays in the lag phage for their kinetics.*

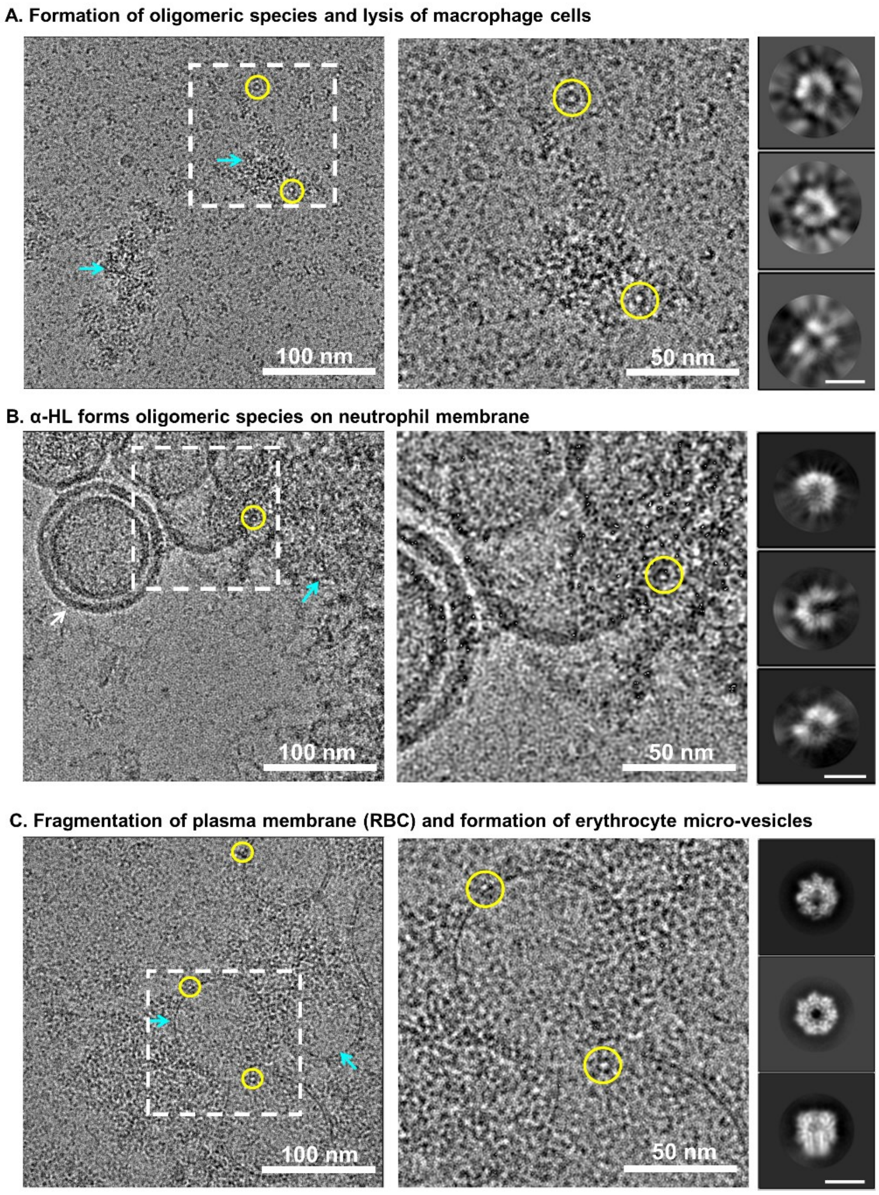


***Figure S7. Formation of oligomeric intermediates and lysis of cellular plasma membrane upon toxin encounter.***

***A-C****. The representative cryo-EM micrograph and an enlarged view (white box) contained toxin-treated fragmented and lysed intracellular debris of RAW264.7* ***(A)****, HL-60 cells* ***(B)****, and RBCs* ***(C)*** *as shown using cyan arrow. Oligomeric species of α-HL shown in yellow circle. The enlarged view showed oligomeric species over the cell surface. The reference-free 2D class averages of the respective oligomeric species identified from different cellular environments shown in the right side. Scale bar 10nm.*

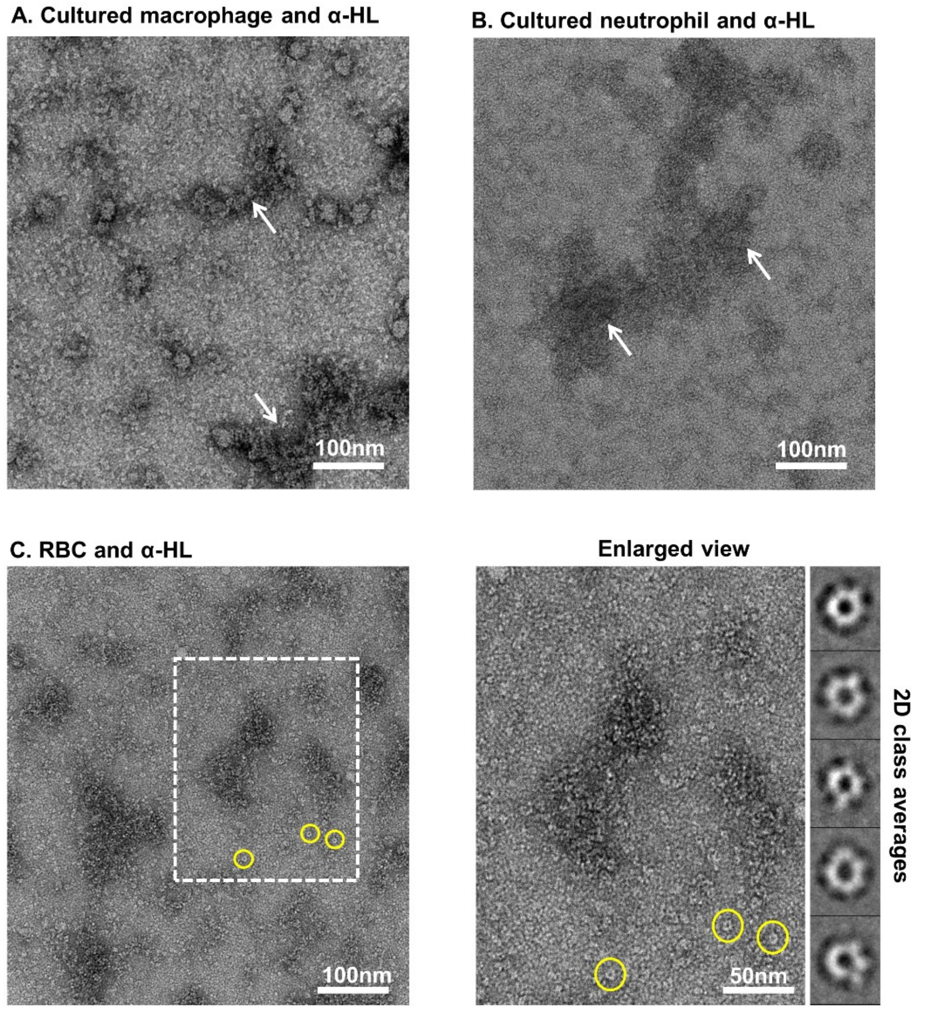


***Figure S8: Cryo-EM sample preparation optimization for native cellular conditions.***

***A-B.*** *NS-TEM image of α-HL treated macrophage (RAW264.7) and neutrophil (HL-60) cells suggested to contain a significant amount of lysed intra-cellular debris (white arrowhead) with very few detectable oligomeric particles.* ***C.*** *NS-TEM images α-HL treated RBC contained sufficient oligomeric toxin (yellow encircled) with reduced cellular debris that was prioritized for an optimal native cellular condition for cryo-EM freezing. Furthermore, 2D class averages showed oligomeric species identified from toxin-treated RBCs.*

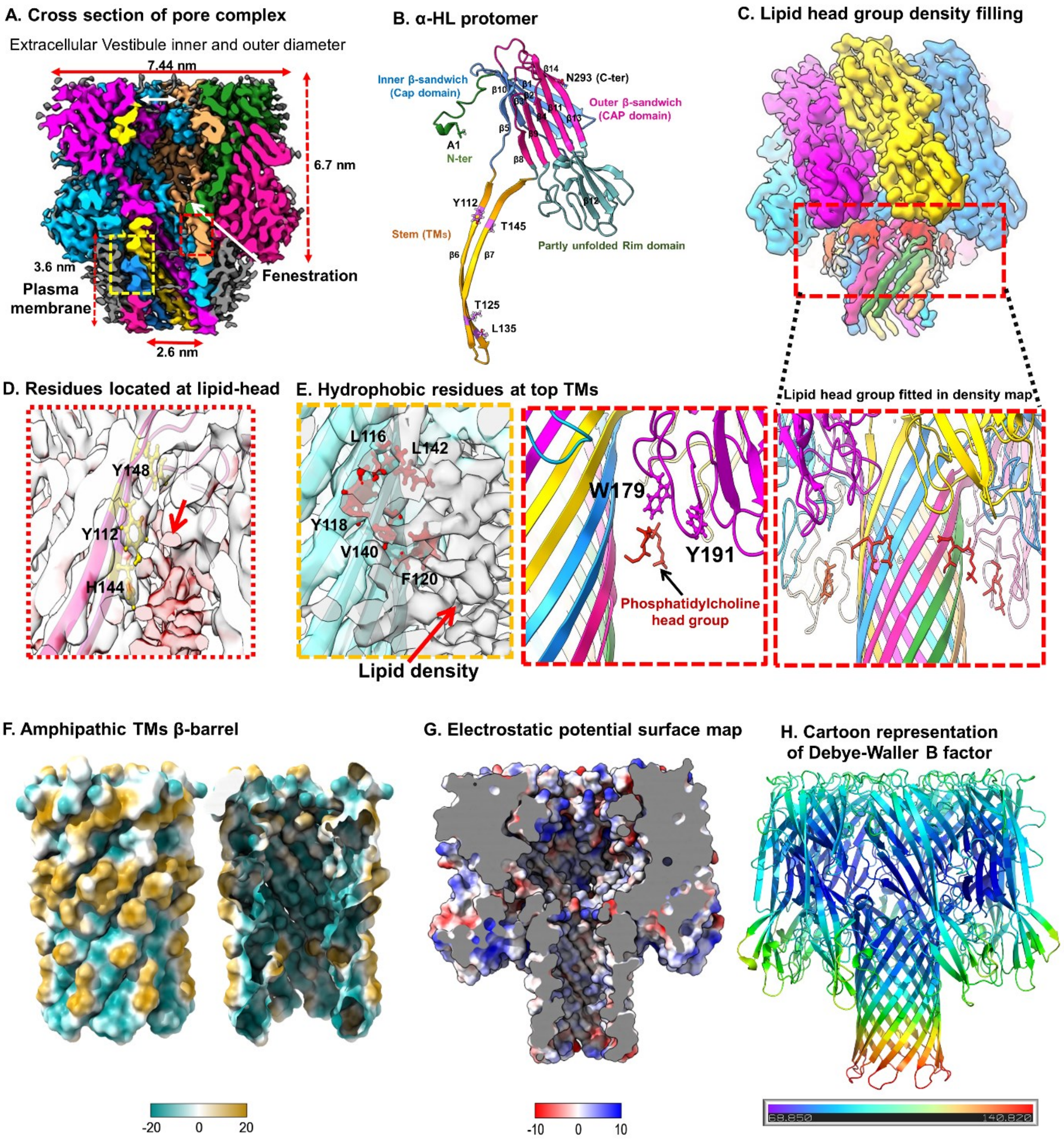


***Figure S9: Structural overviews of heptameric α-HL pore complex and protein-plasma membrane (lipid) interfaces.***

***A.*** *Cross-sectional view of cryo-EM map of α-HL heptameric pore complex displaying the RBC plasma membrane stabilized TMs parts and several other extracellular vestibular regions of the pore complex.* ***B.*** *The distinct segmental analysis of the atomic model of individual α-HL protomer of heptameric pore states.* ***C.*** *The selected enlarged section of the cartoon representation of the atomic model pore complex showing the fitting of the atomic model (ball and stick) of the lipid head group into cryo-EM map located at upper TMs part of pore complex.* ***D.*** *The amphipathic aromatic residues Y112 (from β6) and Y148, H144 (from β7) located at upper TMs (shown in yellow; ball and stick) remained contiguous to fuzzy unmodeled lipid density (red).* ***E****. Large hydrophobic residues (L116, Y118, F120, V140, and Y148) located at upper TMs covered by plasma membrane lipid densities (white in cryo-EM map).* ***F.*** *The hydropathic surface distribution of TMs pore predicted localization of hydrophobic side chain at upper TMs and polar residues in lower TMs (left). While the cross-sectional of inner TMs pore was predominantly polar. Scale bar reflected the relative change in hydrophobic and polar distribution.* ***G.*** *Surface charge distribution of the inner cross-sectional part of the entire channel.* ***H.*** *The relative thermal fluctuations in pore complex.*

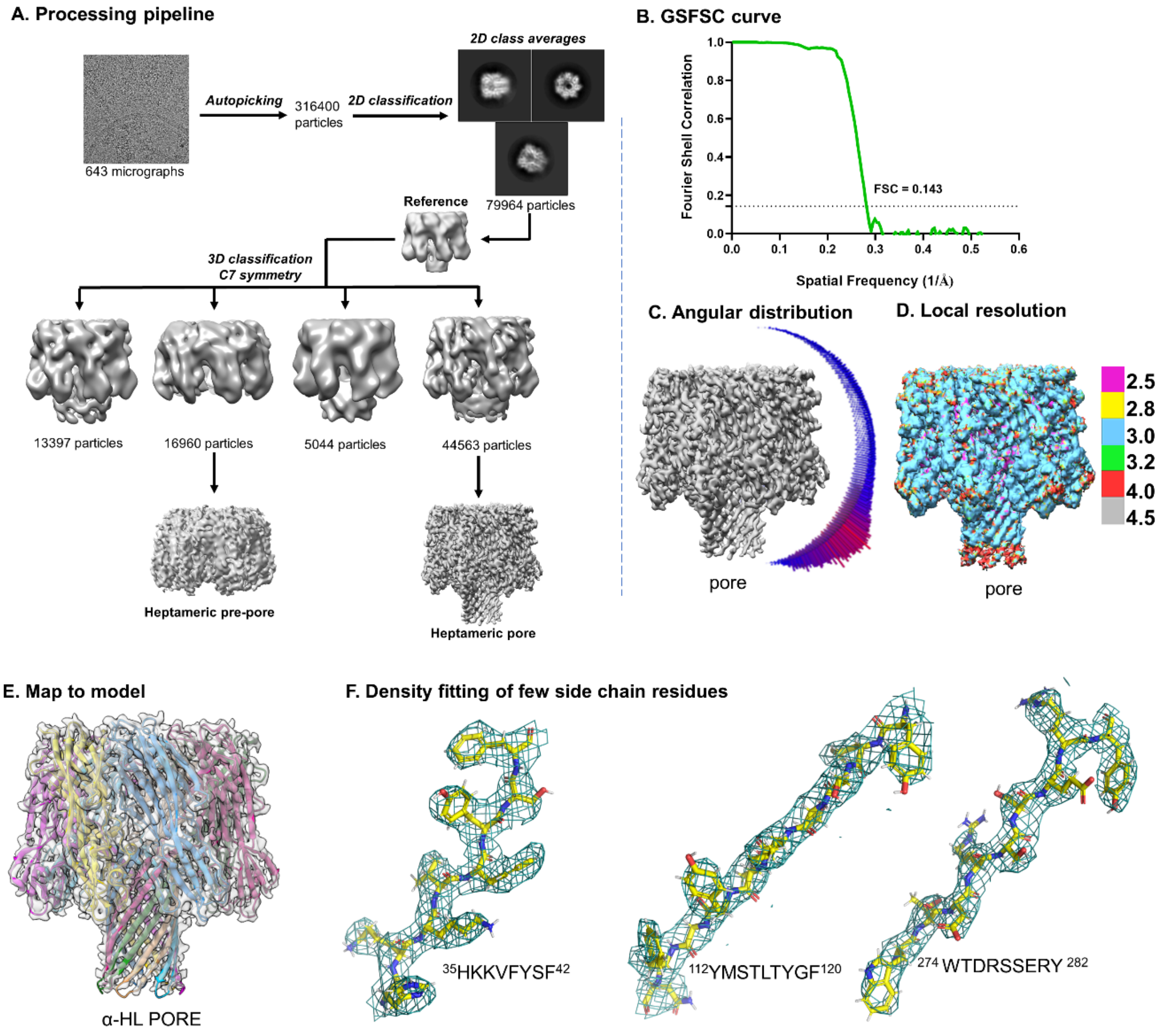


***Figure S10: Cryo-EM data processing pipeline and validation of heptameric species formed in the presence of erythrocyte environment.***

***A.*** *Cryo-EM image processing pipeline for the oligomeric states of α-HL from RBC plasma membrane.* ***B.*** *Gold Standard Fourier Shell Correlation from the two half-maps of heptameric pore states calculated a 3.5 Å global resolution of the refined cryo-EM map of the pore complex.* ***C.*** *An anisotropic distribution of particles in map.* ***D****. A predicted local resolution of 2.5-4Å for the cryo-EM map of heptameric pore states.* ***E.*** *The density of the respective EM map (black) was fitted with an atomic model of the heptameric pore. Each protomer is shown in different colour.* ***F.*** *Fitting of atomic co-ordinates (yellow; ball and stick) with electron density map (shown in green colour mesh) from a few representative segments of α-HL.*

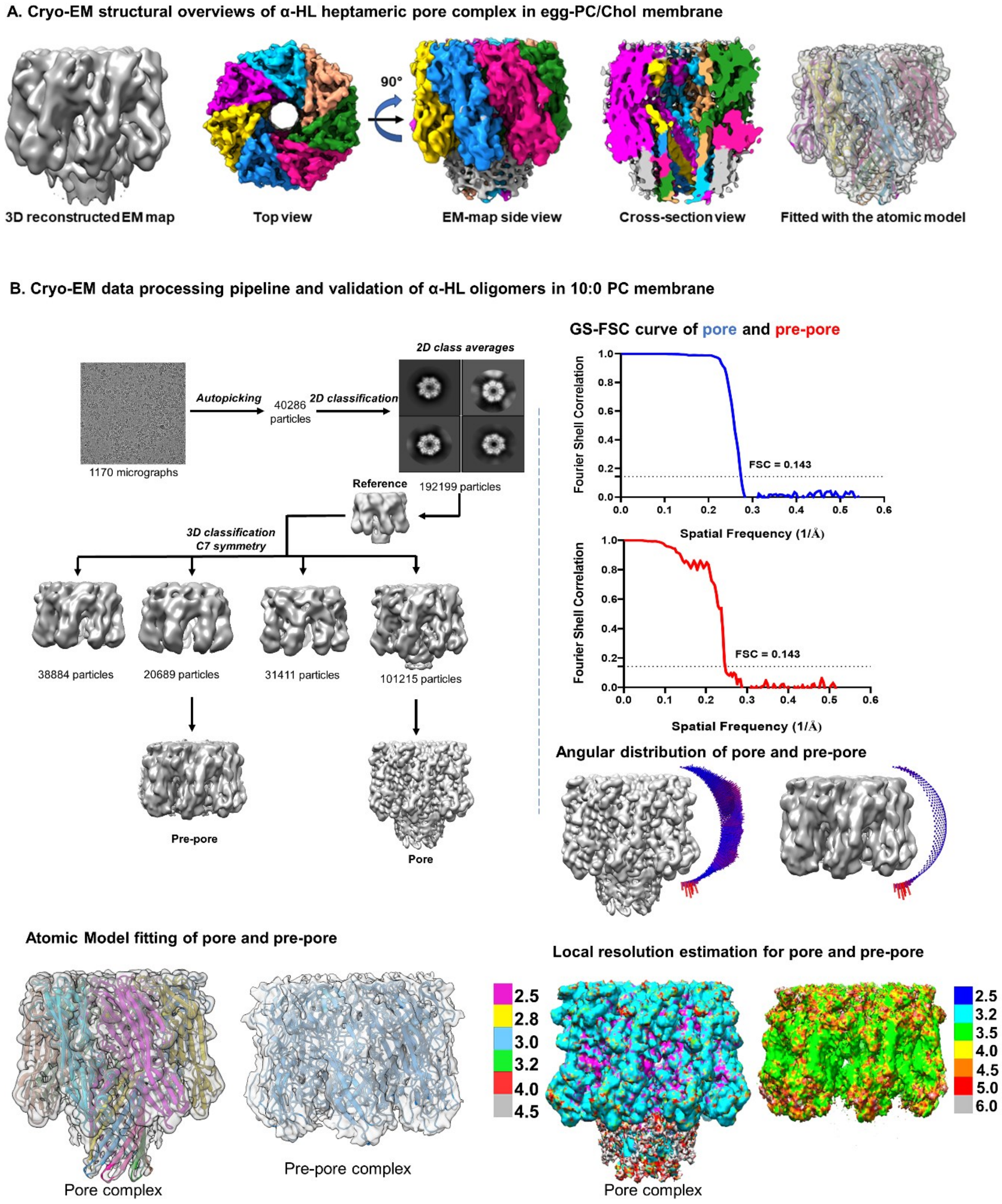


**Figure S11.** ***A.*** *Structural (Cryo-EM) overviews of heptameric pore complex of α-HL isolated from egg-PC/Chol model bio-membranes environments.* ***B.*** *Cryo-EM data processing pipeline for the oligomeric states of α-HL identified from 10:0 PC LUVs. Gold Standard FSC of heptameric pore and pre-pore cryo-EM map had a 3.6 Å and 4.0 Å global resolution respectively. An angular distribution and the local resolution distributions for both the cryo-EM maps of heptameric pore and pre-pore states were depicted. The electron density of the EM map was fitted with their respective atomic models (cartoon representations).*

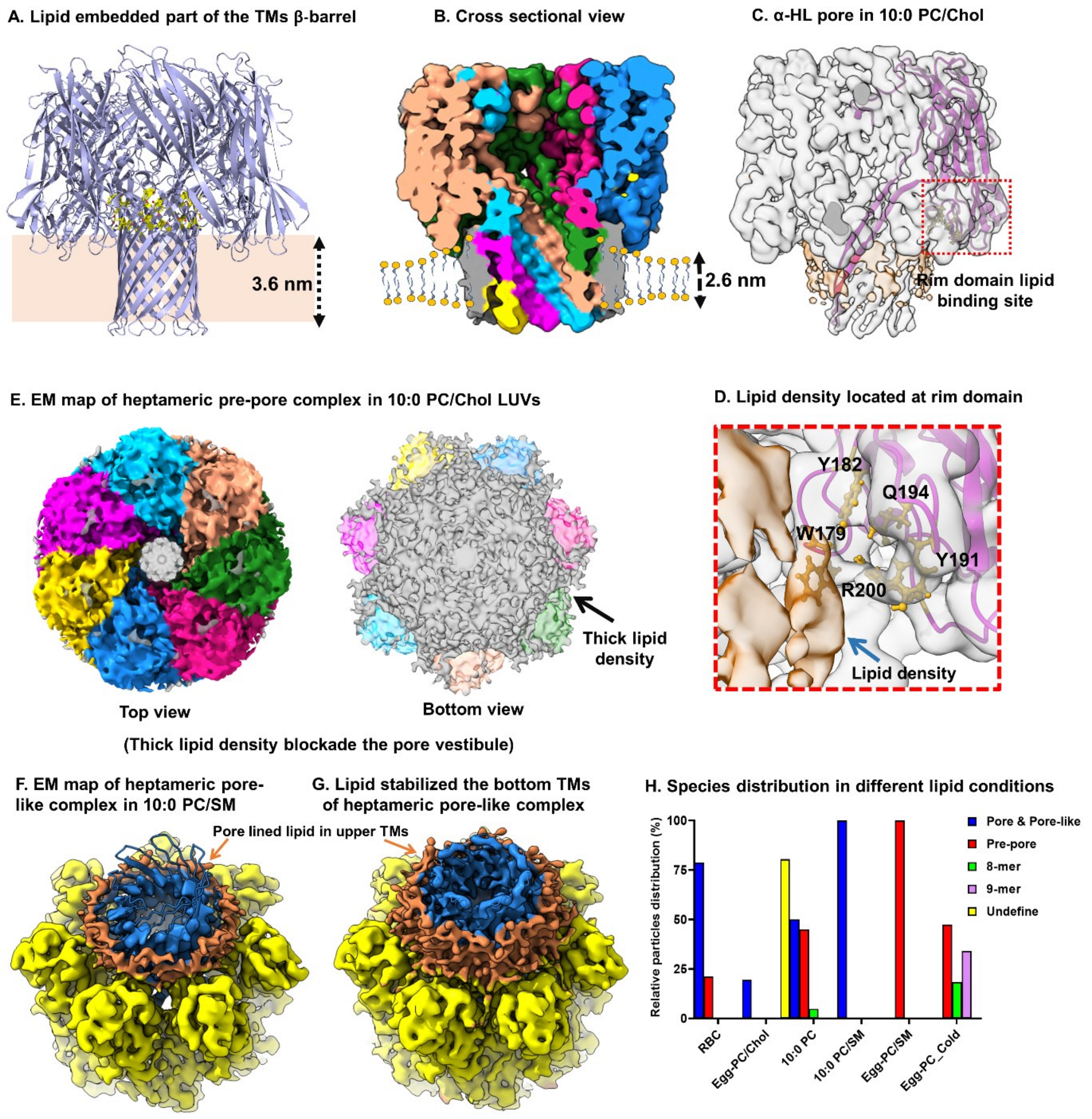


***Figure S12. Lipid re-modulation at protein-lipid interfaces and lipid stabilization of TMs region of α-HL pore and pre-pore complex.***

***A.*** *The schematic representation of α-HL pore structure embedded with a hypothetical 3.6 nm lipid bilayer (light green slab).* ***B.*** *The cross-sectional view of the pore complex (isolated from 10:0 PC LUVs) coated with thick ~3.6nm lipid density (grey) at TMs pore while lipid bi-layer thickness of 10:0 PC LUVs is 2.6nm.* ***C-D.*** *The bottom part of the rim domain showed the plausible lipid-binding residues (W179, Y182, Y191, Q194, and R200) located at the rim domain of α-HL.* ***E.*** *The top and bottom views of heptameric pre-pore complex. A thick lipid density (white) completely blocked TMs pore vestibule.* ***F, G.*** *The cryo-EM map of pore-like states fitted with atomic model identified from 10:0PC/SM lipid membrane at bottom TMs from cryo-EM map. The re-appearance of this missing density at lower TMs with higher map threshold (0.00592). TMs barrel was shown to be coated with a pore line lipid layer (orange).* ***H.*** *The distribution of different oligomeric states of α-HL complex identified from RBC plasma membrane and several model proteo-liposomal membranes.*

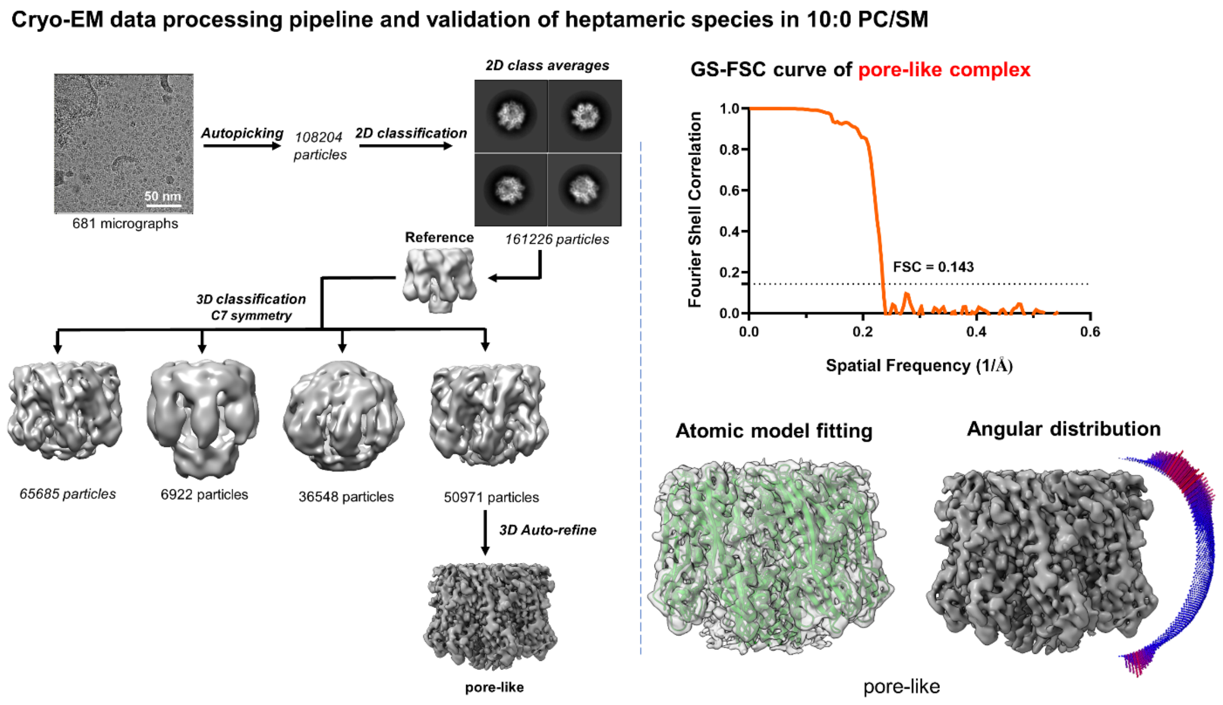


*Figure S13. Cryo-EM data processing pipeline for the oligomeric pore-like states of α-HL generated from 10:0 PC/SM LUVs.*

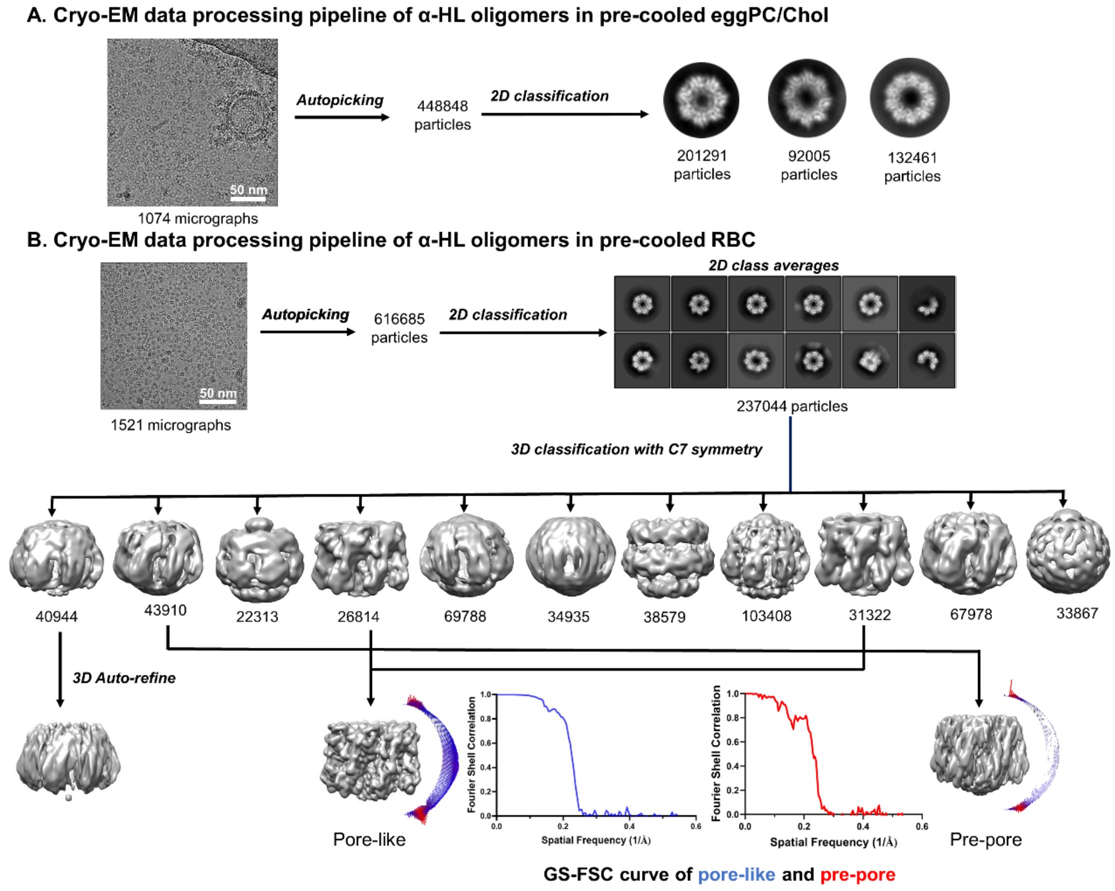


***Figure S14.*** ***A.*** *Cryo-EM data processing pipeline for α-HL incubated with pre-cooled eggPC/Chol liposomes revealed the existence of heptameric, octameric, and nonameric geometrical states.* ***B.*** *A detailed cryo-EM data processing path for cryo-EM structural characterization of heptameric pore-like, pre-pore and twisted oligomeric species of α-HL as predicted from toxin treated pre-cooled RBC membranes. GSFSC of heptameric pore-like and pre-pore cryo-EM map suggested 4.2 Å and 4.0 Å global resolution for both states respectively.*

**Supplemental Information**

**Table S1**: Cryo-EM Data collection, image processing and refinement for membrane and α-HL complex:

| **Data Collection and Processing** | **RBC-pore** | **10:0 PC** | **10:0/SM** | **Pre-cooled RBC** |
|---|---|---|---|---|
| Magnification | 54,000 X | 54,000 X | 54,000 X | 54,000 X |
| Voltage | 200 kV | 200 kV | 200 kV | 200 kV |
| Electron exposure ($e^-$/Å$^2$) | 50 | 50 | 50 | 50 |
| Defocus range (μm) | -1.25 to -2.75 | -1.25 to -2.75 | -1.25 to -2.75 | -1.25 to -2.75 |
| Pixel size (Å) | 0.92 | 0.92 | 0.92 | 0.92 |
| Symmetry Imposed | C7 | C7 | C7 | C7 |
| Number of micrographs | 643 | 1170 | 681 | 3018 |
| Number of particles | 44563 | 101215, 20689 | 50971 | 58136,111858 |
| Map Resolution (Å) | 3.5 | 3.6, 4 | 4.1 | 4.1, 4.0 |
| FSC threshold | 0.143 | 0.143 | 0.143 | 0.143 |
| Map Resolution Range | 3-4 | 3-4, 3.5-5 | - | - |
| Map Sharpening B Factor (Å$^2$) | 100 | 50, 100 | 100 | 100 |
| Oligomer conformation | Pore | Pore and pre-pore | Pore-like | Pore-like, pre-pore |

**Table S2**: Cryo-EM map and model validation for membrane and α-HL complex:

| | | α-HL structural analysis with different lipid bilayer | | | | |
|---|---|---|---|---|---|---|
| **Validation** | **RBC (Pore)** | **10:0 (Pore)** | **10:0 (Pre-pore)** | **10:0/SM (Pore-like)** | **Pre-cooled RBC (Pore-like)** | **Pre-cooled RBC (Pre-pore)** |
| MolProbity Score | 1.55 | 1.60 | 3.59 | 1.70 | 1.62 | 3.60 |
| Clash Score | 4.59 | 5.29 | 69.67 | 7.05 | 5.50 | 30.58 |
| Rotamer Outliers (%) | 0.72 | 0.50 | 8.80 | 1.07 | 0.86 | 12.97 |
| Ramachandran Plot | | | | | | |
| Outlier (%) | 0.10 | 0.15 | 0.05 | 0.12 | 0.16 | 0.12 |
| Allowed (%) | 4.52 | 4.42 | 10.27 | 4.09 | 4.55 | 23.62 |
| Favored (%) | 95.39 | 95.43 | 89.68 | 95.80 | 95.30 | 76.26 |

| Primer Name | Primer Sequence |
|---|---|
| **HL_4RES_FR** | 5'-GGAATTCCATATGATTAATATCAAGACCGGCACCACC-3' |
| **HL_9RES_FP** | 5'-GGAATTCCATATGGCACCACCGATATTGGCAG-3' |
| **HL_TER_RP** | 5'- CCGCTCGAGATTGGTCATTTCCTCTTTTTCCC -3' |
| **HL_278C_FP** | 5'-GATAAATGGACCGATCGTTGTAGCGAACGTTATAAAATTG-3' |
| **HL_278C_RP** | 5'-CAATTTTATAACGTTCGCTACAACGATCGGTCCATTTATC-3' |

| REAGENT | SOURCE | IDENTIFIER |
|---|---|---|
| **Bacterial and virus strains** | | |
| *Escherichia coli* BL21(DE3) | Novagen | Cat# 69450-3CN |
| **Chemicals, peptides, and recombinant proteins** | | |
| **Tris Base** | HiMedia | Cat# TC072M |
| **Sodium Chloride** | Qualigens | Cat# Q15918 |
| **Potassium Chloride** | Merck | Cat# P3911 |
| **Potassium di-hydrogen orthophosphate** | Qualigens | Cat# Q19465 |
| **Di-sodium hydrogen orthophosphate** | SD Fine Chemicals | Cat# S40158 K05 |
| **Luria Broth** | HiMedia | Cat# M575 |
| **Isopropyl ß-D-1-thiogalactopyranoside** | Merck | Cat# I6758 |
| **Imidazole** | Sigma Aldrich | Cat# 792527-500G |
| **2-Methyl-2,4-pentanediol** | Sigma Aldrich | Cat# 8208190100 |
| **L-α-phosphatidylcholine (Egg, Chicken)** | Avanti Polar Lipids | Cat# 840051P-200mg |
| **Cholesterol** | Merck Millipore | Cat# C8997-5G |
| **Ni-NTA Agarose** | Qiagen | Cat# 30210 |
| **Coomassie Brilliant blue R 250** | Merck | Cat# 1.12553 |
| **Precision Plus Protein™ Dual Color Standards** | Bio-Rad | Cat# #1610374 |
| **Amicon Ultra-4 Centrifugal filter unit (10K)** | Merck | Cat# UFC801024 |
| **PC Membranes 0.2µm** | Avanti Polar Lipids | Cat# 610006-1EA |
| **10mm Filter Supports** | Avanti Polar Lipids | Cat# 610014-1EA |
| **Extruder Set with Holder/Heating Block** | Avanti Polar Lipids | Cat# 610000-1EA |
| **Syringe (1000 µL)** | Avanti Polar Lipids | Cat# 610017-1EA |
| **DSPE-PEG (2000) Biotin** | Avanti Polar Lipids | Cat#880129P-10mg |
| **Brain SM** | Avanti Polar Lipids | Cat#860062P-25mg |
| **10:0 PC** | Avanti Polar Lipids | Cat#853025P-25mg |
| **14:0 PC (DMPC)** | Avanti Polar Lipids | Cat#850345C-500mg |
| **16:0 PC (DPPC)** | Avanti Polar Lipids | Cat#850355C-500mg |
| **18:1(Δ9-Cis) PC (DOPC)** | Avanti Polar Lipids | Cat#850375P-1g |
| **Rhodamine B** | Sigma-Aldrich | Cat#83689-1G |
| **Nile-Red** | Sigma-Aldrich | Cat#72485-100MG |
| **4ME 16:0 NBD PE (NBD-DPhPE)** | Avanti Polar Lipids | Cat#810142C-1mg |
| **Streptavidin** | | |
| **Egg Liss Rhod PE** | Avanti Polar Lipids | Cat#810146C-5mg |
| **HL-60 cells** | ATCC | Cat#CRL-3306 |
| **RPMI 1640** | Gibco/Thermofisher Scientific | Cat#11875093 |
| **Foetal Bovine Serum** | Gibco/Thermofisher Scientific | Cat#16000044 |
| **Egg PC** | Avanti Polar Lipids | Cat#131601P-5g |
| **L-Glutamine** | Gibco/Thermofisher Scientific | Cat#25030081 |
| **Dulbecco's Phosphate Buffer Saline** | Sigma-Aldrich | Cat#D5652-10X1L |
| **Bovine Serum Albumin** | Sigma-Aldrich | Cat#A7906 |
| **Apoptosis and Necrosis Quantification Kit Plus (50 assays)** | Biotium | Cat#30065 |
| **BD FACSAria III instrument** | BD Biosciences | N/A |
| **Uranyl Acetate 98%, ACS Reagent** | Polysciences, Inc | Cat# 21447-25 |
| **Superdex 200 Increase 10/300 GL** | Cytiva | Cat# GE28-9909-44 |
| **EM grid (Carbon film 300 mesh, Copper)** | Electron Microscopy Sciences | Cat# CF300-CU |
| **EM grid (Quantifoil R 1.2/1.3 300 Mesh, Copper)** | Electron Microscopy Sciences | Cat# Q3100CR1.3 |
| **Quantifoil R 1.2/1.3, UT, 300 Mesh, Copper** | Electron Microscopy Sciences | Cat# Q3100CR1.3-2nm |
| **4',6-diamidino-2-phenylindole** | Thermofisher Scientific | Cat#D1306 |

| REAGENT | SOURCE | IDENTIFIER |
|---|---|---|
| **Atto 647N Maleimide** | Sigma Aldrich | Cat#05316 |
| **Streptavidin** | Sisco Research Laboratories | Cat#87610 |
